# In search of a core cellular network with single cell transcriptome data in the fruit fly

**DOI:** 10.1101/2021.09.19.460857

**Authors:** Ming Yang, Benjamin R. Harrison, Daniel E.L. Promislow

## Abstract

Along with specialized functions, cells of multicellular organisms also perform essential functions. Whether diverse cells do this by using the same set of genes, interacting in a fixed coordinated fashion, remains a central question in biology. Single-cell RNA-sequencing (scRNA-seq) measures gene expression of individual cells, enabling researchers to discover gene expression patterns that contribute to the diversity of cell functions. Current efforts focus primarily on identifying differentially expressed genes. However, patterns of co-expression between genes are more indicative of biological processes than are the expression of individual genes. Here, we constructed cell type-specific gene co-expression networks using the fly brain transcriptome atlas to search for a core cellular network. We detected a set of highly coordinated genes preserved across cell types and defined this set as the core network. This network was further validated through two additional datasets profiling fly brains or heads. This core has a small size and shows dense connectivity. Modules within this core are enriched for basic cellular functions and gene members of these modules have distinct evolutionary signatures. Overall, we demonstrated that a core cellular network exists in diverse cell types of fly brains and this core exhibits unique topological, structural, functional and evolutionary properties.

## Introduction

Life on Earth has gone through many transitions in organizational complexity (Smith and Szathmary 1997). Among these, the evolution of multicellularity stands out as a key milestone. This transition has occurred independently multiple times across the tree of life and paved the way for tremendous phenotypic expansion and biological diversification (Parfrey and Lahr 2013). Although this has led to the evolution of cell type-specific regulatory pathways that define cells with vastly different functions, all cells in multicellular organisms also carry out common functions that are essential for cell survival. Whether these common functions are supported by a shared network of coexpressed genes, remains a central question in biology (Lim et al. 2013; Hart and Alon 2013). In particular, do all cells utilize the same set of genes in a coordinated fashion to accomplish common functions, to form a core regulatory network?

Cellular phenomena can be characterized by different endophenotypic domains or levels of biological organization, such as the genome, epigenome, transcriptome, proteome, etc. Investigating core functions from these different levels not only provides insight into essential functions of cellular life, but also helps to reveal the evolutionary forces acting at different levels of biological organization (Sorrells and Johnson 2015; Ghadie et al. 2018; Wagner 2012). To identify core functions at each level, researchers have used various strategies such as identifying genes that are expressed constitutively over temporal or spatial scales, and across environments. These genes are typically referred to as ‘housekeeping genes’ and are thought to perform essential functions. Housekeeping genes tend to be evolutionarily ancient (Zhu et al. 2008), highly conserved (Zhang and Li 2004), and are enriched for several functions, including metabolism, RNA binding, protein degradation and cytoskeleton functions (Zhang and Li 2004; Lehner and Fraser 2004). Using somewhat circular logic, core functions are often described based on housekeeping genes. However, we recognize that genes do not work in isolation, but work with each other to carry out biological processes. Co-expression in particular is an indicator of functional relationships (Hughes et al. 2000), and so individual molecular abundances alone may not adequately capture biological organization.

High-throughput methods that generate high-dimensional ‘omic’ data have greatly increased our understanding of molecular and cellular function and organization, in particular through the analysis of molecular networks based on co-expression (Barabasi and Oltvai 2004; Proulx et al. 2005; Thompson et al. 2015; Promislow 2005). Networks consist of nodes connected to one another by edges. In the search for the underlying network structure of cells, researchers have explored many different kinds of edges, including but not limited to gene co-expression, protein-protein interactions (PPI), interactions among transcription factors (TF), TF chromatin occupancy, miRNA-target gene interactions, metabolite covariation, and metabolic reactions (Mitra et al. 2013). Many studies have mainly focused on tissue-specific networks (Greene et al. 2015; Sonawane et al. 2017), and some have also examined networks that show some level of conservation. For example, co-expression network analysis of human and *Arabidopsis* bulk transcriptome data has found a substantial number of gene pairs whose co-expression spans multiple datasets (Lee et al. 2004; He and Maslov 2016). In both analyses, gene pairs expressed across samples were enriched for translation, DNA replication, and regulation of transcription functions, all generally considered to be core cellular functions. Additionally, Skinnider et al. (2021) constructed tissue-specific PPI networks using co-immunoprecipitation within each of seven mouse tissues (Skinnider et al. 2021). They discovered core cellular modules, present in all mouse tissues, composed of evolutionarily ancient proteins, which contrasts with evolutionarily novel accessory modules that are found within individual tissues.

A major drawback of most previous studies investigating core functions from a network perspective is that the networks were inferred from bulk data, which profiles heterogeneous cell populations of an organism, organ, or tissue. Bulk samples face two main limitations for network construction. First, differences in cellular compositions between samples may confound covariation analysis (Farahbod and Pavlidis 2020). Second, measurements that are averaged over thousands of cells in bulk samples make it difficult to detect interactions between genes in individual cells, such as the presence of co-expression patterns and the cell-specificity of these interactions. The compendium of housekeeping genes, initially characterized based on the consistency of their expression, may need revision based on analyses of gene-gene relationships. Housekeeping genes may show clear evidence of co-expression in all cell types; or alternatively, interactions among housekeeping genes may be relatively weak or even cell type-specific. To distinguish these possibilities requires that we define gene networks at a cellular level.

With the advent of single-cell RNA sequencing (scRNA-seq), we have an unprecedented opportunity to reveal gene relationships in specific cellular contexts and to probe cellular-level networks (Trapnell 2015; Tanay and Regev 2017). One recent study used scRNA-seq from mouse brain samples to construct gene co-expression networks, comparing the topology of networks built from different levels of cell type hierarchy (i.e., from broad to specific classes) (Harris et al. 2021). Their results show well-preserved gene-gene relationships at each level, and suggest the existence of a core co-regulatory network in the brain. However, they did not directly compare cellular networks across cell types, which leaves the possibility that what appear as core networks at more integrated levels may not manifest when examined by cell type, or by individual cell. Considering the state of the field and the considerable amount of scRNA-seq data now available, we are led to ask several fundamental questions. First, can we identify shared co-expression patterns between pairs of genes across different cell types; Second, how common are these shared co-expressed gene pairs among cell types? Lastly, do these shared co-expressed genes point to a core cellular network, and if so, what properties does this core network manifest?

To investigate these questions, we used a published scRNA-seq dataset derived from whole fly brains (Davie et al. 2018), and constructed cell type-specific gene co-expression networks. We then looked for shared edges across cell types. We found a population of gene-pairs that are more common across cell types than expected by chance, and yet, none of these edges were found in all cell types. We then determined that those edges that were common among most but not all cell types tended to be near, but below, the correlation threshold we used to construct gene co-expression networks in the remaining cells, suggesting that the edges of a core network show consistent high and varying co-expression strength over cell types. The network we uncovered is small relative to the number of commonly expressed genes, which indicates that core cellular functions are either more limited than previous analysis suggests, or that gene co-expression is not a salient feature of functions common to all cells. We further described the core network’s topological properties, functional enrichment, and the evolutionary age of its constituent genes. We validated our findings in two independent datasets, one based on cells from the fly brain, and a second from the fly head, demonstrating the extensibility of our analytical approach. To our knowledge, this is the first study of a core cellular network among cell types using single-cell transcriptome data in the fly brain and marks a shift in paradigm for investigating commonalities in a molecular endophenotypic domain from a network perspective at the single-cell level.

## Results

### Construction of cell cluster-specific gene co-expression networks

We analyzed published fly brain atlas data obtained from whole-cell fly brain samples of female and male *Drosophila* using 10X Genomics library preparation methods (Davie et al. 2018). The original dataset contained expression profiles of 17,473 genes in 56,902 high-quality brain cells. The cells were grouped into 116 cell clusters, 74 of which were annotated to known fly brain cell types (Davie et al. 2018). Here we built cell cluster-specific gene co-expression networks in females and males separately, and we investigated the consistency of covarying gene pairs across these cell clusters. We selected 68 cell clusters that contained at least 200 cells and filtered expressed genes in each cell cluster individually (**Methods**). The number of genes expressed per cell cluster ranged from 2,683 in the Tm5ab cell cluster of females, to 6,727 in cell cluster 0 of males (**Fig. S1**). In total, there were 7,795 genes expressed in at least one cell cluster, 2,088 of which were expressed in all 68 cell clusters (**Fig. S1**). scRNA-seq data are typically sparse, with the expression of many genes falling below the limits of detection. Sparsity, measured as the percentage of zeros in the data for the 2,088 commonly expressed genes, was below 50% for all but one cell cluster (cluster 32 in males, **Supplementary Material, section 1)**. After excluding this cell cluster from further analysis, the remaining cell clusters had less than 48% sparsity. Throughout, we focus on the 2,088 commonly expressed genes to identify covarying gene pairs and co-expression networks, within and across these 67 cell clusters.

An overview of our pipeline is shown in **Figure 1**. We used the bigScale2 algorithm (Iacono et al. 2019) to compute cell cluster-specific gene correlation matrices and then built cell cluster-specific gene co-expression networks by thresholding these gene correlation matrices with a top percentile-based cutoff approach (**Method and Fig. S2**). While there is no objective criterion for choosing a gene correlation threshold, we employed a signal-to-noise score approach, which indicates the consistency of networks constructed using subsets of original data at a given threshold, to guide the threshold value selection (**Supplementary Material, section 1**). The results showed that most cell cluster networks had the highest signal-to-noise score when constructed at the top 5% threshold (**Supplementary Material, section 1, Fig. S3 and S4)**, hence we applied this cutoff and transformed gene correlation matrices into gene co-expression networks (**Fig. S5)**. At this threshold we identified 1,942,694 co-expressed gene pairs (89.16% of all possible pairs among 2,088 genes) that occurred in at least one of the 67 cell cluster-specific networks.

**Figure 1.**
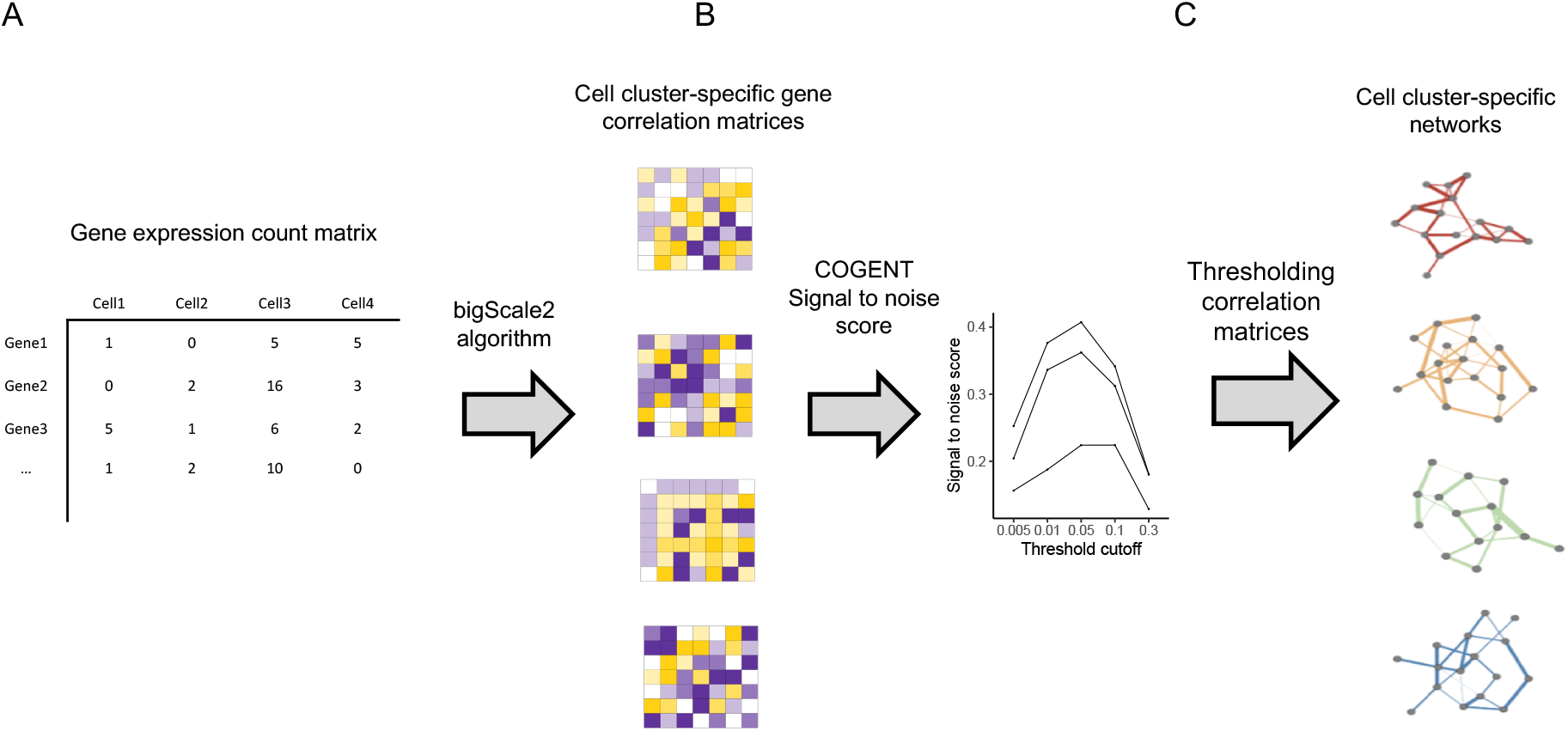
Pipeline overview of cell cluster-specific gene co-expression network construction. The pipeline starts with a gene expression count matrix (A) and computes a gene correlation matrix for each cell cluster using the bigScale2 algorithm (B, Iacono et al. 2019). Signal-to-noise scores are used to evaluate network robustness at different percentile thresholding values for each absolute gene correlation matrix individually, out of which a global optimal thresholding value is selected (C). Applying the selected thresholding cutoff value, gene correlation matrices are transformed into gene co-expression networks.

### Co-expression networks in fly brain cell clusters are highly context-dependent

If a core cellular network exists, we expect its edges to be present in all cell clusters. We define the number of cell clusters in which each edge (i.e., each co-expressed gene pair) is detected as that edge’s ‘commonality’. The distribution of commonality scores was right skewed, with more than 75% of the edges specific to fewer than five cell clusters and only 0.4% of edges common to more than 30 cell clusters (**Fig. 2A**). The largest recorded commonality score was 64, observed for only two edges. Given that 67 cell clusters were analyzed, no edges were found to be present in every cluster. As a complement to the observed edge commonality distribution, we also plotted the gene commonality distribution, where gene commonality indicates the number of cell clusters in which a given gene shared an edge with at least one other gene. The gene commonality distribution showed that most genes had one or more edges in the majority of cell clusters, and 613 genes had at least one edge in all 67 cell clusters (**Fig. 2A**). Thus, commonly expressed genes were frequently co-expressed with other genes, though the specific co-expression partners vary among different cell clusters.

**Figure 2.**
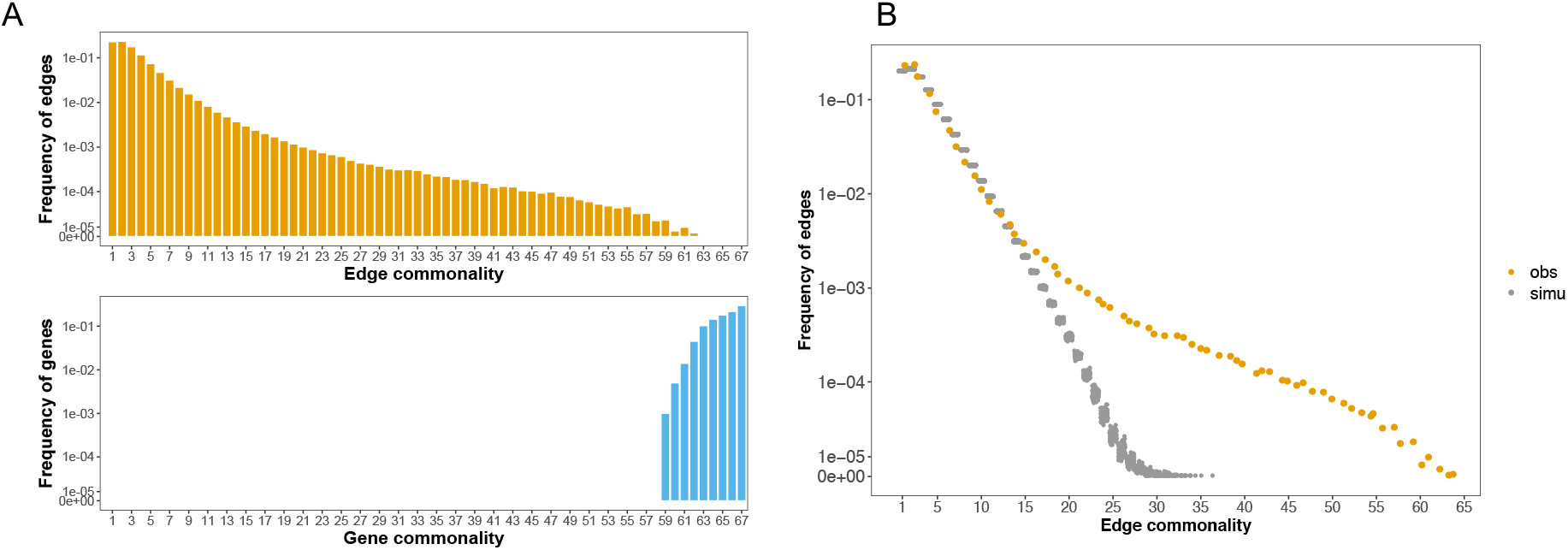
Cell cluster-specific co-expression networks share many more edges than random networks. A. Edge and gene commonality distributions. The commonality of an edge indicates the number of cell clusters in which that edge is detected (top). The commonality of a gene refers to the number of cell clusters in which that gene is detected as being co-expressed with one or more other one gene (bottom). The y-axis shows the frequency of genes or edges in the corresponding commonality score group. B. The observed edge commonality distribution (yellow) compared with the null expectation derived from network randomization (gray). Network randomization was performed 100 times for each cell cluster individually with network size (number of nodes and edges) and gene degree (number of co-expressed gene partners per gene) fixed.

### Recurrently co-expressed genes in multiple cell clusters

The fact that no edges were found to occur in all cells suggests either that a core cellular network does not exist, or that perhaps our method for detecting co-expression was unable to identify all edges of a core network. To determine if our method uncovers gene pairs that recurrently co-express in multiple cell clusters, we next asked to what extent the observed edge commonality distribution differed from the null expectation, where gene co-expression occurs randomly among commonly expressed genes. We evaluated this in two ways. First, we derived a mathematical expectation for the probabilities of edge commonality using the binomial distribution (**Methods**). This calculation shows that most gene pairs were expected to co-express in only a few cell clusters. For example, for a gene pair to be co-expressed in exactly 2, 3 or 4 cell clusters, the probability values were 0.1970, 0.2247, or 0.1892, respectively, and the probability became smaller than 0.0001 for cell cluster ≥14 (**Methods**). In a null model based on random gene co-expression, the probability of one gene pair not co-expressed in any of the 67 cell types is 0.0321. Thus, we would expect to find 2,108,888 unique gene pairs to occur in one or more clusters, among the 2,088 commonly expressed genes (**Methods**). This number is larger than the observed 1,942,694 gene pairs, indicating that some genes recurrently co-express in multiple cell clusters. A full comparison of this analytically predicted distribution and the observed edge commonality showed that the two distributions agreed well at lower, more cell-specific commonality. However, we observed a clear excess of shared edges relative to the frequency expected for those expressed in 7 or more clusters **(Fig. S6**). Second, we compared the deviation between the observed edge commonality distribution and a null distribution sampled using network randomization (**Methods**). This comparison showed that the observed distribution was enriched for high commonality edges. For instance, none of the randomizations generated an edge commonality larger than 36, while the observed distribution included hundreds of such edges (**Fig. 2B**), further supporting our observation that many edges occur in more cell clusters than we would expect by chance. This pattern is robust to the percentile cutoff values used in network construction (**Fig. S7**). Additionally, using a loose co-expression threshold that incorporates more edges in each cell cluster network led to edge commonality distributions that more closely resembled random networks (**Fig S7**). These results therefore suggest that there exists a set of co-varying genes that occur more repeatedly than expected by chance across diverse cellular contexts, pointing to a core cellular network composed of genes that are co-expressed across cell types.

### Rank aggregation analysis reveals a core cellular network with sub-threshold edges

An ideal core network would consist of edges that appear in every cell cluster. However, in practice, the identification of gene co-expression edges based on gene expression data involves the risk of false positives and false negatives. In the data sets we analyzed, no edges were present in every surveyed cell cluster. This could be due to our parameter value choices, such as the correlation matrix thresholding cutoff, or alternatively, these edges or gene pairs could truly be cell cluster specific, and they are not co-expressed in all cell clusters. We hypothesized that if a core network exists but certain edges are missing as false negatives, these edge members should be consistently highly ranked among the gene pairs in all cell clusters, even if they are ranked below the significance threshold in some cell types. To test this possibility, we performed a rank aggregation analysis for network edges in those cell clusters where they do not pass the threshold for defining an edge. Specifically, we extracted edges from a given edge commonality group *k* (they belong to the top 5% in *k* cell clusters) with *k* ranging from 1 to 51, covering the spectrum of edge commonalities scores, and examined each edge’s ranking vector in the remaining *67-k* cell clusters. The rank aggregation algorithm estimated a P value per edge ranging from 0 to 1, with a small value indicating an edge is ranked consistently higher across cell clusters, and a larger value meaning an edge’s rank distribution over cell clusters follows a random pattern.

To enable fair comparison between edge commonality groups, we randomly sampled 100 edges in each group and computed their rank aggregation scores. The enrichment of highly ranked edges indeed increases with edge commonality, a strong indication that edges in high commonality groups are consistently highly ranked among the gene pairs in those cell clusters, despite being below the threshold cutoff in the remaining cell clusters (**Fig. 3A**). In particular, more than 95 percent of edges sampled from the edge commonality groups ≥ 43 were highly ranked in the remaining cell clusters (P < 0.05), whereas fewer than 1% of the tested edges shared by 7 or fewer cell types (edge commonality ≤ 7) reached the 5% significance threshold in other clusters. Based on this rank aggregation analysis and a P value < 0.05 cutoff, we chose edge commonality 43 to define core network edges, which resulted in a core network in the Drosophila brain with 205 genes connected by 2,140 edges (**Table S1**).

**Figure 3.**
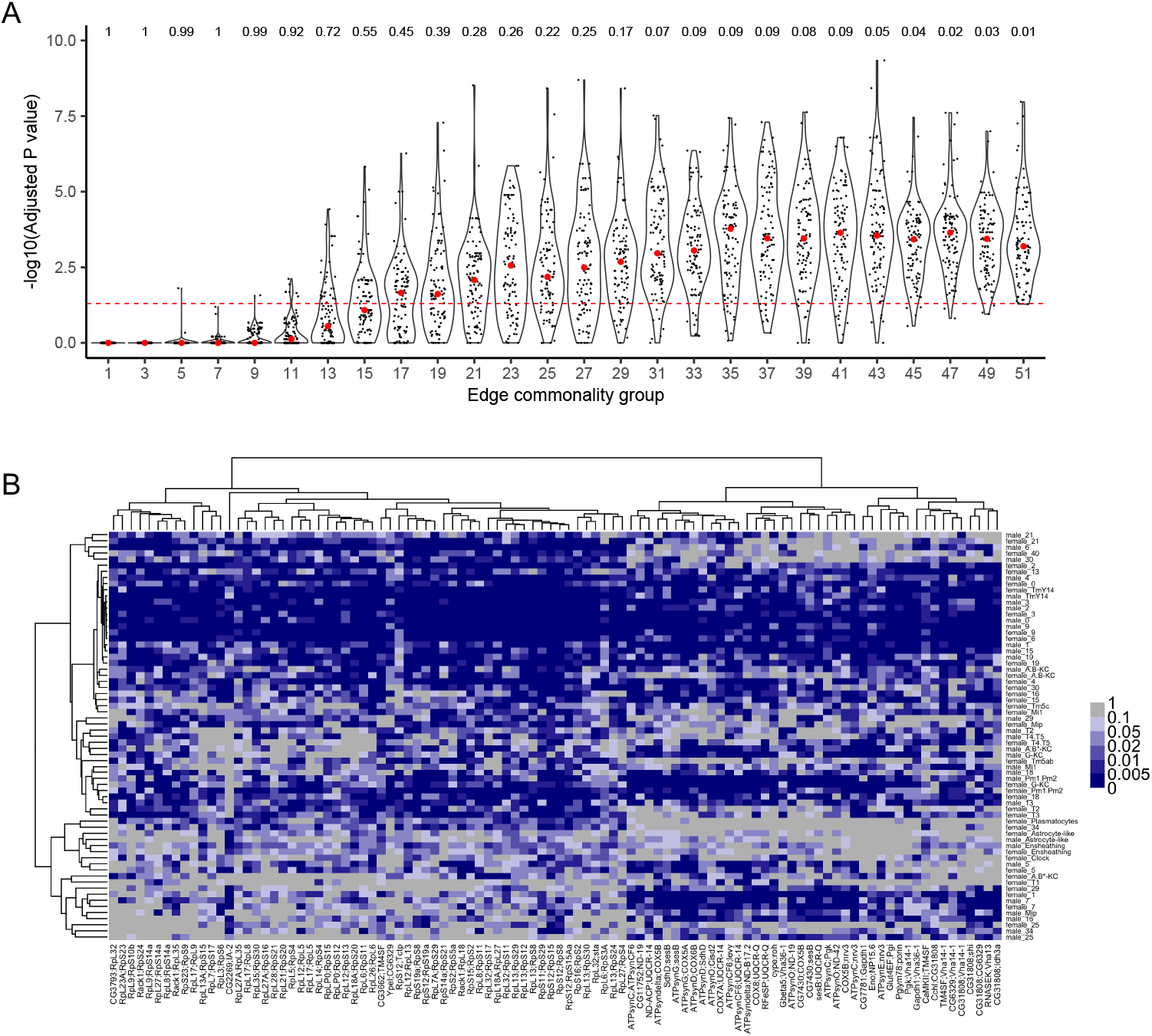
Identifying a core using an edge commonality cutoff based on rank aggregation analysis. A. Adjusted P-value distributions of 100 sampled edges in different edge commonality groups using a rank aggregation method. Edge commonality groups were selected to cover the full range of edge commonality scores. Each violin plot illustrates the distribution of adjusted P values for the sampled edges for each edge commonality group. Each dot represents an edge, with red dots indicating median values. The dashed red line indicates an adjusted P value of 0.05. The number above each violin indicates the percentage of edges with adjusted P values smaller than 0.05 in the corresponding edge commonality group. B. Heatmap of the normalized ranks for the 100 sampled edges with a commonality score of 43 across cell clusters. Each edge has a rank *r* in each cell cluster based on its absolute correlation value, from which a normalized rank r’ = (r-1)/(R-1) is calculated, where R is the total number of edges in a cell network. A smaller normalized rank value indicates this edge is highly ranked among gene pairs and is encoded as dark blue colors in the heatmap. The row labels are cell cluster annotations and the column labels are gene pairs. The trees on both sides are generated via hierarchical clustering using the ‘hclust’ function in R with the “ward.D2” method on the Euclidean-based distances.

While we define the core network based on edges found in at least 43 cell clusters, it is possible that that these edges occur repeatedly but are limited to the same set of cell types, and so perhaps they may have more cell-specificity than we might expect for a hypothetical core network, whose structure should be nearly the same regardless of cell type. To explore this possibility, we clustered cell types based on the rank of 100 randomly sampled edges from edge commonality group 43 (**Fig. 3B**). This analysis revealed a group of about 20 cell clusters that contain nearly all of the core edges. The remaining cell clusters contained a variety of network edges along with highly ranked, but sub-threshold, network edges in their co-expression matrices. This indicates that the relative strength of the edges in the core varies by cell cluster.

Of all *Drosophila* organs, the brain may contain the greatest diversity of cell types that are currently resolved by scRNA-seq studies (Li et al. 2021). Having identified a network that is shared across cell types of the *Drosophila* brain, we then validated our analysis in an independent study of the same organ (Baker et al. 2021) (**Supplementary Material, section 2**), and extended our analysis to a study of cells from whole fly heads, which allowed us to survey an even wider range of cell types (Li et al. 2021) (**Supplementary Material, section 3)**. The independent study of the *Drosophila* brain included samples of adult flies of both sexes, treated with either sucrose, or sucrose supplemented with cocaine, and profiled by scRNA-seq (Baker et al. 2021). We focused our analysis on sucrose samples for both sexes, and found 1,738 commonly expressed genes across 47 cell clusters. Following network construction as above, we identified a network of 323 edges and 88 genes (**Fig. S8**). As in the analysis described above, in this brain core network, we did not identify edges found in all cell types. However, edges found in most cell types ranked higher than expected by chance in the remaining cell types (**Fig. S8**).

Turning to the analysis of the entire fly head, among the 76 cell clusters identified, we found 842 commonly expressed genes, among which we identified a network of 21 edges and 20 genes (**Fig. S9**). Compared to the fly brain data, the fly head data included at least nine additional cell clusters, which may represent cell types outside of the brain, and opens the possibility of evaluating the persistence of the putative core network across a wider range of cell types. Here again, the head core network did not have edges in all cells, but those expressed in most cells ranked highly in the remaining cell types (**Fig. S9**). We next asked if there was a set of genes observed commonly across all three network analyses. In fact, we found 62 genes that were shared by the core network from fly brain atlas data and the second brain study (dataset of Baker et al. 2021) and seven genes shared with the fly head study (dataset of Li et al. 2021). Compared to random samples from the commonly expressed genes in each data set, these overlaps among the real networks were highly significant (empirical P value = 0.00099, calculated as the proportion of random samples whose overlapped gene numbers were larger than the observed one, **Fig. S8H and S9H**).

### Topological, functional and evolutionary signatures of the core cellular network

Having defined a core cellular network, we next examined its topological, functional and evolutionary properties. To evaluate the topological properties of the core network, we calculated its clustering coefficient, and compared it to an ensemble of coefficients from pseudo-core networks each with the same number of genes, edges, and degree distribution as the observed one. A high clustering coefficient indicates that nodes of a network are densely connected to each other. The observed clustering coefficient value, which has a range of 0 to 1, was 0.68, much higher than the mean simulated value of 0.13 (range 0.11 - 0.16) (**Fig. S10**), suggesting the core network has a dense connection structure.

Many complex gene networks can be divided into modules, where genes are more highly interconnected within modules than between modules (Newman 2003). Modules identified from gene co-expression networks tend to take part in the same biological processes or pathways (Ruprecht et al. 2017; Wolfe et al. 2005). To explore the structural and functional organization of this core, we decomposed it into highly connected modules using the Markov Clustering Algorithm (**Methods**). In total, we identified four modules, each with at least five gene members (**Fig. 4A and 4B; Table S2**). Most gene pairs within modules were positively correlated, while the between-module connections tended to be negative (**Fig. S11)**. We then annotated each module’s biological function through Gene Ontology (GO) enrichment analysis. The results revealed an array of housekeeping functions enriched within different modules (**Fig. 4C; Table S3**), including pathways associated with ATP metabolism, glycolysis, translation, and proteostasis. The largest module (module 3) contained 83 genes, and was enriched for ATP metabolic functions, which suggests highly coordinated expression of genes involved in energy production. The second largest module (module 2) contained 70 genes and was enriched for ribosome related functions, such as cytoplasmic translation, suggesting tight correlation of expression among genes encoding ribosomal proteins across cells. We note that this analysis was limited to mRNA, and so this module does not contain the stoichiometrically synthesized rRNAs. Module 4 had 29 genes associated with glycolysis, a process central to cellular energy homeostasis. Module 1 formed a fully interconnected subnetwork without any edges to other modules (**Fig. 4B)**, and its gene members were highly enriched for heat-shock proteins (HSP) or co-chaperones, key players in protein folding.

**Figure 4.**
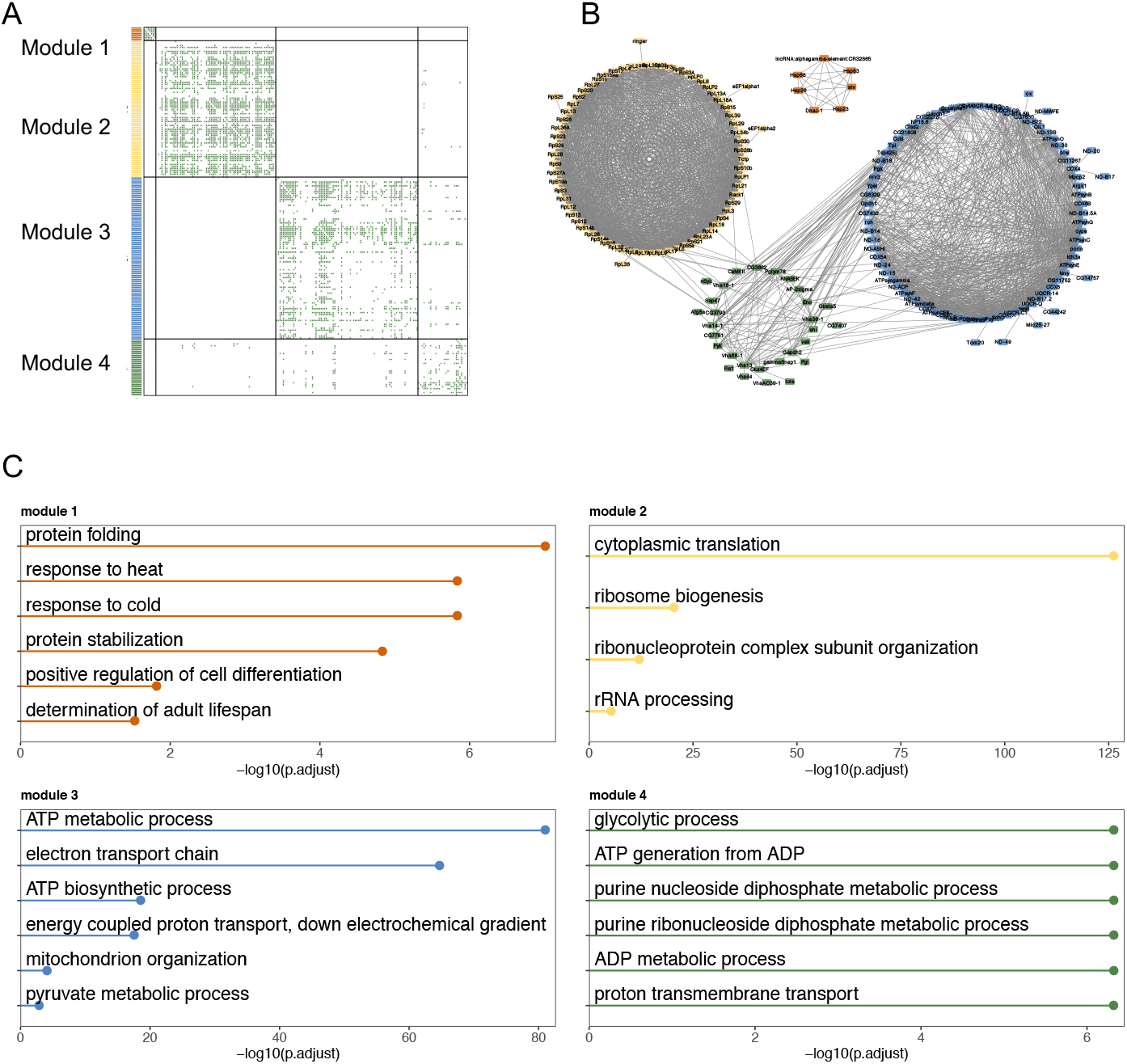
Gene modules in the defined core network. A. Heatmap of gene co-expression relationships and decomposed modules in the defined core cellular network. Modules that have at least five gene members are highlighted in different colors and numerically indexed. B. Network visualization of the core modules. C. Enriched GO terms for each core module. We used the R package ‘clusterProfiler’ to perform gene set enrichment analysis of Gene Ontology with a Bonferroni correction and an adjusted P value cutoff of 0.05. In each module, the top terms are shown (up to 6). A full list of enriched GO terms for each module is provided in **Table S3**.

To characterize the evolutionary signature of each module, we used phylostratigraphy, assigning each gene in each module to one of 10 different evolutionary time periods (Domazet-Lošo et al. 2017) (**Table S4**). Broadly speaking, genes in the core network were enriched for genes with ancient origins, compared with other commonly expressed genes in the fly brain (**Fig. S12A**). Looking in detail, the four modules of the core network differed significantly with respect to the evolutionary age of their constituent genes (**Fig. S12B**). Gene members of the ribosomal (module 2) and protein folding (module 1) modules predated the divergence of the eukaryota, while those of the ATP metabolic (module 3) and glycolysis (module 4) modules included genes distributed across both ancient and relatively recent evolutionary time periods. While this analysis measures the age of the gene members at the network nodes, rather than the age of network edges, these diverse age signatures of different core modules suggest that some modules arose by integration of both young and old genes, perhaps involving step-wise recruitment of young genes into ancestral core modules.

## Discussion

To what extent do all cells in an organism rely on a common core of interacting genes? Would such a network be detectable among the expression patterns of diverse cell types? To investigate these questions, we examined cell type-specific gene co-expression networks using fly brain scRNA-seq and fly head snRNA-seq data. We describe an approach to find shared components among multiple co-expression networks, identifying a small and highly clustered core network. We validate this result, finding structurally and functionally similar networks in two independent datasets, one from the fly brain, and a second from fly heads, the latter of which includes a greater diversity of cell types. The core network we identify is composed of four co-expression modules, each of which has distinct functional enrichment and distribution of gene age.

While the search for core networks is not new (Almaas et al. 2005; Neph et al. 2012; Skinnider et al. 2021), our study is distinct in at least three ways from previous work. First, instead of relying on consistent expression levels of individual genes to identify genes common across cell types, we examined covariation between genes as the measure of functional commonality. This focus provides not only a stricter criterion to infer gene function (Hughes et al. 2000), but also has the potential to capture conserved gene regulatory networks (Yu et al. 2003; Stuart et al. 2003; Segal et al. 2003). Second, a large body of studies has relied heavily on PPI data to derive biological networks and find commonalities (Liu et al. 2020; Skinnider et al. 2021; Huttlin et al. 2021). While these studies are informative, they suffer from the bias that PPI data often lack information on the degree of cell specificity of such interactions, and are enriched for highly-studied proteins, which may give an incomplete picture of network structure (Skinnider et al. 2018; Gillis et al. 2014; Schaefer et al. 2015). In our study, we analyzed transcriptome data, which interrogate almost all genes in the genome and are less biased with respect to knowledge from prior databases or existing literature. Third, we identified covarying gene pairs using scRNA-seq data, which unlike bulk transcriptome data, or broad-scale PPI data, can be defined by cell type, even within a single biological sample. In contrast to bulk transcriptomic analysis and PPI data, where the cellular specificity of each interaction is largely ambiguous, scRNA-seq enabled us to build cell type-specific networks at a resolution that was previously impossible.

### Topological properties of the core network

With our current parameter choices, the defined core network is remarkably small when compared to the much larger network of cell-specific interactions. In particular, in the fly brain, the relative core size, calculated as the 2,140 core edges divided by 108,942, the number of edges in a cell cluster-specific network, is only 2%. This number is relatively small compared with previous studies of different biological networks. For example, Skinnider et al. (2021) constructed tissue-specific PPI networks for seven mouse tissues and found that those universal PPIs shared by all tissues typically occupy 3.7% of the total number of PPIs in a tissue (Skinnider et al. 2021). Neph et al. (2012) built TF interaction networks for 41 cell types in humans and found that 5% of interacting TFs were common to all cell types (Neph et al. 2012). Almaas et al. (2005) used flux-balance analysis to study active metabolic reactions of *Escherichia coli* in 30,000 diverse simulated environments and predicted that 90 of 758 (11.9%) reactions were always active (Almaas et al. 2005).

Another prominent feature of this core network is its dense connectivity. The gene network architecture we observed, which embodies extensive cell-type specific interactions along with a shared and densely connected core, echoes findings from other types of biological networks. For example, Liu et al. (2020) identified 13,764 PPIs in yeast across nine environments and found that 60% of PPIs were found in only one environment (Liu et al. 2020). They also show that PPIs that were present in eight or more environments formed ‘tight’ modules of high node degree, while PPIs present in three or fewer environments formed less-connected modules of smaller node degree. Protein interaction networks based on just two human cell lines revealed that shared interactions tend to reside in dense subnetworks and correspond to known protein complexes such as the exosome and the COP9 signalosome (Huttlin et al. 2021). Similarly, network analyses of gene co-expression from bulk transcriptomics in *Arabidopsis* or in humans suggest a highly connected core, which appears alongside an extensive number of condition-specific gene interactions (He and Maslov 2016; Lee et al. 2004). Taken together, these results suggest a universal organizing principle in biological systems, where widely shared components of interaction networks are relatively small and densely connected (Milo et al. 2002, 2004; Csermely et al. 2013).

### Is there a core network active in all cells?

To determine whether there is a core co-expression network common to all cells, we focused our analysis on genes that were expressed across all cell clusters. This enabled us to directly compare the diverse co-expression patterns among a common set of genes. Single-cell sequencing data suffer from a high level of sparsity. It has been estimated that the proportion of zeros in bulk RNA-seq data is usually 10% - 40%, while in scRNA-seq data it can be as high as 90% (Jiang et al. 2022). Such sparsity may be due to both technical reasons, such as the lack of efficiency in capturing the single-cell transcriptome during library preparation, and biological reasons, such as the stochastic nature of gene expression (Sarkar and Stephens 2021). Data sparsity, as a universal analytical challenge for scRNA-seq data, is an active research area, and little consensus has been reached on how to handle this challenge (Lähnemann et al. 2020). Our approach involved deriving cell cluster-specific co-expression matrices and only then did we look at covariation of each gene pair across cell clusters, and we found that the patterns of co-expression networks that we detected were robust to relatively high levels of sparsity (**Supplementary Material, section 1**).

Even among the genes expressed in all cell clusters of the fly brain, we found that most gene co-expression was cell cluster-specific, and yet there also existed a significantly large number of genes whose co-expression occurred among multiple cell clusters. This enrichment of shared co-expressed genes suggests the existence of a core co-expression network. While we identified edges common to many cell clusters, there was no edge that was shared by all cell clusters. We consider two alternative explanations for this observation.

First, it is possible that a common core co-expression network for all brain cells does not exist, and that cell co-expression networks are so diverse as to lack such rigid network structure across the cells of the *Drosophila* brain. Alternatively, while sparsity does not appear to present a problem for our analysis, there is still the possibility that a core does exist, but that the statistical inference of gene co-expression and the threshold values we use to build the network might not fully resolve the core in all cells. To define the core, we specified multiple parameters, including a gene correlation metric, a correlation threshold to select co-expressed genes, and an edge commonality cutoff to extract a core network. Given that our definition of gene interactions is based on statistical inference of correlations, we might simply fail to observe a real interaction in one or more cell clusters due to type II error (false negatives). As we explored this parameter space, moving from stringent to relaxed parameter values, a few features of the network became apparent. In particular, more relaxed parameter values revealed co-expressed gene pairs observed in all cell clusters in the brain, but this inherently increased the risk of type I error (false positive gene pairs), as revealed by network permutation. While the network that we define lacks edges shared by all cell clusters, edges that are present in most clusters tend to be highly ranked in those cell types where they do not reach the threshold of significance. Thus, their absence from the top 5% of gene pairs in some cells may be partly explained by their relatively weak co-expression strengths compared to the cell cluster-specific gene pairs. Together, our analysis indicates the existence of a core cellular network, though the size and composition that we define is conditional on parameter choices.

It is likely that other factors not examined here also contribute to the size, structure and/or function of a core network. These could include cell/organ type, sex, genotype, age, and so forth. For example, one recent study using bulk RNA-seq data to examine gene co-expression network dynamics with age in bats identified a small core network whose size decreases with age (Bernard et al. 2022). That said, we find that the basic structure of our defined core network occurs in both sexes when measured in the fly brain, a complex organ with highly heterogeneous cell type composition. Projecting such an analysis to more organs, or even to a whole fly, would very likely reveal an even smaller core, as the inclusion of a larger set of diverse cells would lead to a smaller set of universal genes, and universal edges. Indeed, in our analysis of the fly head, which contains more cell types than the fly brain, the uncovered core network was indeed smaller, yet still significantly intersecting in gene identity with the core network from a unique sample of fly brains. We therefore speculate that the relative core size that we observe in the fly brain data, which includes 2% of cell cluster-specific network edges, might be an overestimate of the true core network size for all cells in *Drosophila*. Future studies should apply our approach on an organism-wide scale, in search of a small but ubiquitous core network testing found in all cells in an organism.

### Limitation and future directions

In this study, we sought a core of interacting genes found across cell clusters in the fly brain. While the work described here benefits from access to high quality single-cell transcriptome data, there are still several caveats worth noting. First, the fly brain cell atlas (Davie et al. 2018) was generated using a mixture of two genotypes, and with cells from individuals of several ages. Thus, genotype or age-specific gene co-expression patterns are yet to be revealed. Future studies targeting individual genotypes and/or specific age groups might uncover a dynamic picture of how core networks change over age or genotype conditions. Second, we inferred co-expressed gene pairs from gene expression data statistically. Gene co-expression is not proof of gene co-regulation, which may be more indicative of functional relationships. Further experimental work is needed to validate the functional implications of these gene pairs. Lastly, we caution that the analysis on fly brain samples is based on scRNA-seq data, and the fly head samples were processed using single nuclear RNA-seq. These two approaches might capture different aspects of cell activity (Wu et al. 2019; Denisenko et al. 2020; Thrupp et al. 2020).

Our approach, which is generic and interpretable, can be extended to incorporate more data, such as additional tissues, species, or biological conditions in future studies. Although current single cell techniques yield data with high levels of sparsity, somewhat ambiguous cell type resolution, and other challenges, we anticipate that similar and more complete data are on the horizon. We expect that more comprehensive knowledge of gene expression in all cells may reveal that a core co-expression network, shared by all cells under all conditions, may be very limited in size, perhaps even non-existent. This possibility does not rule out the idea that all cells share common functions that are required for cell survival, but that these functions are not always dependent on gene co-expression, or that the co-expression within such a network is weak relative to the many conditional co-expression relationships that occupy cellular networks.

## Methods

### Dataset collection and preprocessing

We downloaded the fly brain atlas data from NCBI Gene Expression Omnibus (GEO GSE107451). The original dataset contains expression data for 17,473 genes in 56,902 high-quality brain cells grouped into 116 cell clusters. As a quality control step, we first removed 668 cells in a cell cluster named ‘Hsp’, as they represent stressed cells (Davie et al. 2018). We then removed cells that had fewer than 200 expressed genes, fewer than 500 total unique molecular identifier counts, or a total fraction of mitochondrial gene expression exceeding 30%. These criteria led to the removal of another 42 cells, leaving 56,192 cells. These cells were assigned to 115 cell clusters from which we selected 37 cell clusters in females and 31 cell clusters in males that had at least 200 cells each.

We filtered genes for each cell cluster in each sex individually by removing genes that were expressed in less than 15 cells, or in fewer than 0.5% of cells in that cell type. This gene filtering procedure led to 7,795 genes as expressed in at least one cell type, 2,088 of which were commonly expressed in all 67 cell clusters (**Fig. S1**).

### Constructing cell type-specific gene co-expression networks

We used the bigScale2 algorithm (Iacono et al. 2019) to compute a gene-gene correlation matrix for each cell cluster in each sex. This algorithm was tailored to mitigate the impact of sparse counts at the single-cell level. It first groups cells into homogenous cell clusters, then performs differential expression (DE) analysis between all pairs of clusters. With N clusters, we obtain N*(N-1)/2 unique comparisons, and each comparison generates one z-score for each gene, indicating the likelihood of an expression change between the corresponding two clusters. Finally, bigScale2 uses transformed z-scores instead of original expression values to calculate Pearson correlation coefficients. This z-score transformation allows us to detect correlations that would otherwise be missed by drop-out events and other technical artifacts. Example scatter plots of z-scores of gene pairs within cell clusters are presented in **Fig. S2**. In **Section 1** of the **Supplementary Material**, we further demonstrate the ability of this algorithm to provide robust estimates even with sparse data. To select highly correlated gene pairs for inclusion in a gene co-expression network, we employed a signal-to-noise score approach and calculated this score across various top percentile-based threshold values in each cell cluster separately. The highest signal-to-noise score frequently occurs at a threshold value taking the top 5% of edges across cell clusters (**Fig. S3 and S4**). Thus, we ranked gene pairs by their absolute correlation values in each cell cluster separately, and placed the top 5% of correlated gene pairs into a cell cluster-specific co-expression network, with the corresponding absolute correlation values ranging from *r=*0.39 to 0.85.

### Evaluating gene and edge commonality distributions

To evaluate commonality and specificity across cell cluster-specific networks, we plot the node and edge commonality distributions. The commonality of a node (gene) refers to the number of cell clusters in which this gene is found to be co-expressed (edge) with at least one other gene. The commonality of an edge linking a given pair of genes refers to the number of cell clusters in which that specific edge is detected. We derived a mathematical approximation for the probability of a gene pair to be co-expressed in a given number of cell types. As we focused on 2,088 commonly expressed genes and selected the top 5% of highly correlated genes in each cell type, in randomized data, a gene pair would have a probability *p* = 0.05 as being co-expressed in any one cell type. Examining 67 cell clusters and using the binomial distribution, the probability *P*(*k*) of a gene pair being co-expressed in *k* cell types would equal

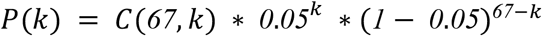

Where *C*(67, *k*) is the combinatorial number describing the number of ways of picking *k* items from a pool of 67 cell clusters (that is, 67 choose *k*, or 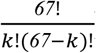). Following this equation, the probability of a gene pair not being co-expressed in any cell type is 0.0321 at *k* = 0. Given the 2,088 commonly expressed genes, 2,108,888 non-redundant gene pairs were expected to co-express in at least one cell type.

### Network randomization

To obtain a null distribution for edge commonality distributions, we used a network randomization approach. We randomized the edges in each cell type-specific network individually, keeping the gene connectivities fixed using the rewire function from the R package iGraph (Csardi and Nepusz 2006). A set of randomizations for all 67 cell clusters resulted in one pseudo edge commonality distribution. We performed the randomization procedure 100 times and used the ensemble of the 100 pseudo edge commonality distributions as the null distribution.

### Rank aggregation analysis

Each gene pair or edge has a rank based on its absolute correlation value in a given cell cluster. To determine if one edge is ranked consistently high across a set of cell clusters based on its absolute correlation value, we used the aggregateRanks function from the R package RobustRankAggreg (Kolde et al. 2012). This function is based on a probabilistic model of order statistics and computes a derived P value for each edge. The derived P value ranges from 0 to 1 and serves as an upper bound of the computationally expensive exact P value, with a small value indicating one edge is ranked consistently higher across cell types. We chose edge commonality groups covering the full range of edge commonality scores, and within each edge commonality group, we randomly sampled 100 edges. For each edge, we first collected the cell clusters in which that edge was absent (i.e., was not in the top 5% of correlations), and then calculated the derived P values from these cell clusters. The derived P values were corrected for multiple testing using the p.adjust function in R with the Benjamini-Hochberg method, referred to as adjusted P values hereafter. We plotted the adjusted P value distribution of the 100 sampled edges for each edge commonality group separately.

### Module decomposition and functional annotation of the core

To decompose the core cellular network into highly connected modules, we used the mcl function from the R package MCL (Jäger 2015), which implements a Markov cluster algorithm to identify clusters in networks. After module detection, we performed Gene Ontology (GO) enrichment analysis of genes in each module using the R package clusterProfiler (Wu et al. 2021), with a Bonferroni correction and an adjusted P value cutoff of 0.05. Significant GO terms were identified and refined to reduce redundant GO terms via the simplify method from the clusterProfiler package.

### Assigning genes into evolutionary age groups

We downloaded data from a previous study to assign genes into different evolutionary age groups using a phylostratigraphy framework (Domazet-Lošo et al. 2017). This framework allows us to date the evolutionary origination time of a gene by identifying its homologs across the tree of life. There were 13,794 genes assigned to 12 age groups in the original publication, 2,002 of which overlapped with the 2,088 expressed genes in this study, including 911 genes in the oldest age group “CellLife”, 641 in “Eukaryota”, 87 in “Opisthokonta”, 122 in “Metazoa”, 32 in “Eumetazoa”, 73 in “Bilateria”, 14 in “Protostomia”, 16 in “Arthropoda”, 13 in “Pancrustacea”, 43 in “Insecta”, 39 in “Diptera” and 11 in the youngest age group “Drosophila”.

### Data access

The fly brain atlas dataset analyzed during the current study is available at GEO with accession number GSE107451. The dataset from Baker et al. (2021) is available at GEO with accession number GSE152495. The dataset from Li et al. (2021) is available from the Fly Cell Atlas website (https://flycellatlas.org/). R scripts for data analyses are available from the following GitHub repository: https://github.com/mingwhy/fly.brain.core_coexpr.net.

## Acknowledgements

This work was supported in part by National Institute on Aging grants R21AG56872901, R01AG057330, and R01AG063371. This work was in part facilitated through the use of advanced computational, storage, and networking infrastructure provided by the Hyak supercomputer system at the University of Washington.

## Author contribution

M.Y. and D.P. designed the study. M.Y. collected and analyzed the data. M.Y., B.H. and D.P. interpreted the data and wrote the manuscript.

## References

Almaas E, Oltvai ZN, Barabási A-L. 2005. The activity reaction core and plasticity of metabolic networks. PLoS computational biology 1: e68. doi: 10.1371/journal.pcbi.0010068

Baker BM, Mokashi SS, Shankar V, Hatfield JS, Hannah RC, Mackay TFC, Anholt RRH. 2021. The Drosophila brain on cocaine at single-cell resolution. Genome research 31: 1927–1937. doi: 10.1101/gr.268037.120

Barabasi A-L, Oltvai ZN. 2004. Network biology: understanding the cell’s functional organization. Nature reviews genetics 5: 101–113. doi: 10.1038/nrg1272

Bernard G, Teulière J, Lopez P, Corel E, Lapointe F-J, Bapteste E. 2022. Aging at evolutionary crossroads: longitudinal gene co-expression network analyses of proximal and ultimate causes of aging in bats. Molecular biology and evolution 39: msab302. doi: 10.1093/molbev/msab302

Csardi G, Nepusz T. 2006. The igraph software package for complex network research. InterJournal, complex systems 1695: 1–9.

Csermely P, London A, Wu L-Y, Uzzi B. 2013. Structure and dynamics of core/periphery networks. Journal of Complex Networks 1: 93–123. doi: 10.1093/comnet/cnt016

Davie K, Janssens J, Koldere D, de Waegeneer M, Pech U, Kreft Ł, Aibar S, Makhzami S, Christiaens V, González-Blas CB. 2018. A single-cell transcriptome atlas of the aging Drosophila brain. Cell 174: 982–998. doi:10.1016/j.cell.2018.05.057

Denisenko E, Guo BB, Jones M, Hou R, de Kock L, Lassmann T, Poppe D, Clément O, Simmons RK, Lister R. 2020. Systematic assessment of tissue dissociation and storage biases in single-cell and single-nucleus RNA-seq workflows. Genome biology 21: 1–25. doi: 10.1186/s13059-020-02048-6

Domazet-Lošo T, Carvunis A-R, Albà M, Šestak MS, Bakarić R, Neme R, Tautz D. 2017. No evidence for phylostratigraphic bias impacting inferences on patterns of gene emergence and evolution. Molecular biology and evolution 34: 843–856. doi: 10.1093/molbev/msw284

Farahbod M, Pavlidis P. 2020. Untangling the effects of cellular composition on coexpression analysis. Genome research 30: 849–859. doi: 10.1101/gr.256735.119

Ghadie MA, Coulombe-Huntington J, Xia Y. 2018. Interactome evolution: Insights from genome-wide analyses of protein–protein interactions. Current opinion in structural biology 50: 42–48. doi: 10.1016/j.sbi.2017.10.012

Gillis J, Ballouz S, Pavlidis P. 2014. Bias tradeoffs in the creation and analysis of protein–protein interaction networks. Journal of proteomics 100: 44–54. doi: 10.1016/j.jprot.2014.01.020

Greene CS, Krishnan A, Wong AK, Ricciotti E, Zelaya RA, Himmelstein DS, Zhang R, Hartmann BM, Zaslavsky E, Sealfon SC. 2015. Understanding multicellular function and disease with human tissue-specific networks. Nature genetics 47: 569–576. doi: 10.1038/ng.3259

Harris BD, Crow M, Fischer S, Gillis J. 2021. Single-cell co-expression analysis reveals that transcriptional modules are shared across cell types in the brain. Cell Systems 12: 748–756. doi: 10.1016/j.cels.2021.04.010

Hart Y, Alon U. 2013. The utility of paradoxical components in biological circuits. Molecular cell 49: 213–221. doi: 10.1016/j.molcel.2013.01.004

He F, Maslov S. 2016. Pan-and core-network analysis of co-expression genes in a model plant. Scientific reports 6: 1–11. doi: 10.1038/srep38956

Hughes TR, Marton MJ, Jones AR, Roberts CJ, Stoughton R, Armour CD, Bennett HA, Coffey E, Dai H, He YD. 2000. Functional discovery via a compendium of expression profiles. Cell 102: 109–126. doi: 10.1016/S0092-8674(00)00015-5

Huttlin EL, Bruckner RJ, Navarrete-Perea J, Cannon JR, Baltier K, Gebreab F, Gygi MP, Thornock A, Zarraga G, Tam S. 2021. Dual proteome-scale networks reveal cell-specific remodeling of the human interactome. Cell 184: 3022–3040. doi: 10.1016/j.cell.2021.04.011

Iacono G, Massoni-Badosa R, Heyn H. 2019. Single-cell transcriptomics unveils gene regulatory network plasticity. Genome biology 20: 1–20. doi: 10.1186/s13059-019-1713-4

Jäger ML. 2015. MCL: Markov Cluster Algorithm. R package version 1.0. https://CRANR-project.org/package=MCL.

Jiang R, Sun T, Song D, Li JJ. 2022. Statistics or biology: the zero-inflation controversy about scRNA-seq data. Genome Biology 23: 1–24. doi: 10.1186/s13059-022-02601-5

Kolde R, Laur S, Adler P, Vilo J. 2012. Robust rank aggregation for gene list integration and meta-analysis. Bioinformatics 28: 573–580. doi: 10.1093/bioinformatics/btr709

Lähnemann D, Köster J, Szczurek E, McCarthy DJ, Hicks SC, Robinson MD, Vallejos CA, Campbell KR, Beerenwinkel N, Mahfouz A, et al. 2020. Eleven grand challenges in single-cell data science. Genome Biology. doi: 10.1186/s13059-020-1926-6

Lee HK, Hsu AK, Sajdak J, Qin J, Pavlidis P. 2004. Coexpression analysis of human genes across many microarray data sets. Genome research 14: 1085–1094. doi: 10.1101/gr.1910904

Lehner B, Fraser AG. 2004. Protein domains enriched in mammalian tissue-specific or widely expressed genes. Trends in Genetics 20: 468–472. doi: 10.1016/j.tig.2004.08.002

Li H, Janssens J, de Waegeneer M, Kolluru SS, Davie K, Gardeux V, Saelens W, David F, Brbic M, Leskovec J. 2021. Fly Cell Atlas: a single-cell transcriptomic atlas of the adult fruit fly. bioRxiv. doi: 10.1101/2021.07.04.451050

Lim WA, Lee CM, Tang C. 2013. Design principles of regulatory networks: searching for the molecular algorithms of the cell. Molecular cell 49: 202–212. doi: 10.1016/j.molcel.2012.12.020

Liu Z, Miller D, Li F, Liu X, Levy SF. 2020. A large accessory protein interactome is rewired across environments. Elife 9: e62365. doi: 10.7554/eLife.62365

Milo R, Itzkovitz S, Kashtan N, Levitt R, Shen-Orr S, Ayzenshtat I, Sheffer M, Alon U. 2004. Superfamilies of evolved and designed networks. Science 303: 1538–1542. doi: 10.1126/science.1089167

Milo R, Shen-Orr S, Itzkovitz S, Kashtan N, Chklovskii D, Alon U. 2002. Network motifs: simple building blocks of complex networks. Science 298: 824–827. doi: 10.1126/science.298.5594.824

Mitra K, Carvunis A-R, Ramesh SK, Ideker T. 2013. Integrative approaches for finding modular structure in biological networks. Nature Reviews Genetics 14: 719–732. doi: 10.1038/nrg3552

Neph S, Stergachis AB, Reynolds A, Sandstrom R, Borenstein E, Stamatoyannopoulos JA. 2012. Circuitry and dynamics of human transcription factor regulatory networks. Cell 150: 1274– 1286. doi: 10.1016/j.cell.2012.04.040

Newman MEJ. 2003. The structure and function of complex networks. SIAM review 45: 167–256. doi: 10.1137/S003614450342480

Parfrey LW, Lahr DJG. 2013. Multicellularity arose several times in the evolution of eukaryotes (Response to DOI 10.1002/bies. 201100187). Bioessays 35: 339–347. doi: 10.1002/bies.201200143

Promislow D. 2005. A regulatory network analysis of phenotypic plasticity in yeast. The American naturalist 165: 515–523. doi: 10.1086/429161

Proulx SR, Promislow DEL, Phillips PC. 2005. Network thinking in ecology and evolution. Trends in ecology & evolution 20: 345–353. doi: 10.1016/j.tree.2005.04.004

Ruprecht C, Proost S, Hernandez-Coronado M, Ortiz-Ramirez C, Lang D, Rensing SA, Becker JD, Vandepoele K, Mutwil M. 2017. Phylogenomic analysis of gene co-expression networks reveals the evolution of functional modules. The Plant Journal 90: 447–465. doi: 10.1111/tpj.13502

Sarkar A, Stephens M. 2021. Separating measurement and expression models clarifies confusion in single-cell RNA sequencing analysis. Nature genetics 53: 770–777. doi: 10.1038/s41588-021-00873-4

Schaefer MH, Serrano L, Andrade-Navarro MA. 2015. Correcting for the study bias associated with protein–protein interaction measurements reveals differences between protein degree distributions from different cancer types. Frontiers in genetics 6: 260. doi: 10.3389/fgene.2015.00260

Segal E, Shapira M, Regev A, Pe’er D, Botstein D, Koller D, Friedman N. 2003. Module networks: identifying regulatory modules and their condition-specific regulators from gene expression data. Nature genetics 34: 166–176. doi: 10.1038/ng1165

Skinnider MA, Scott NE, Prudova A, Kerr CH, Stoynov N, Stacey RG, Chan QWT, Rattray D, Gsponer J, Foster LJ. 2021. An atlas of protein-protein interactions across mouse tissues. Cell 184: 4073–4089. doi: 10.1016/j.cell.2021.06.003

Skinnider MA, Stacey RG, Foster LJ. 2018. Genomic data integration systematically biases interactome mapping. PLoS computational biology 14: e1006474. doi: 10.1371/journal.pcbi.1006474

Sonawane AR, Platig J, Fagny M, Chen C-Y, Paulson JN, Lopes-Ramos CM, DeMeo DL, Quackenbush J, Glass K, Kuijjer ML. 2017. Understanding tissue-specific gene regulation. Cell reports 21: 1077–1088.

Sorrells TR, Johnson AD. 2015. Making sense of transcription networks. Cell 161: 714–723. doi: 10.1016/j.cell.2015.04.014

Stuart JM, Segal E, Koller D, Kim SK. 2003. A gene-coexpression network for global discovery of conserved genetic modules. science 302: 249–255. doi: 10.1126/science.1087447

Tanay A, Regev A. 2017. Scaling single-cell genomics from phenomenology to mechanism. Nature 541: 331–338. doi: 10.1038/nature21350

Thompson D, Regev A, Roy S. 2015. Comparative analysis of gene regulatory networks: from network reconstruction to evolution. Annual review of cell and developmental biology 31: 399– 428. doi: 10.1146/annurev-cellbio-100913-012908

Thrupp N, Frigerio CS, Wolfs L, Skene NG, Fattorelli N, Poovathingal S, Fourne Y, Matthews PM, Theys T, Mancuso R. 2020. Single-nucleus RNA-seq is not suitable for detection of microglial activation genes in humans. Cell reports 32: 108189. doi: 10.1016/j.celrep.2020.108189

Trapnell C. 2015. Defining cell types and states with single-cell genomics. Genome research 25: 1491–1498. doi: 10.1101/gr.190595.115

Wagner A. 2012. Metabolic networks and their evolution. In Evolutionary systems biology, pp. 29–52, Springer. doi: 10.1007/978-1-4614-3567-9_2

Wolfe CJ, Kohane IS, Butte AJ. 2005. Systematic survey reveals general applicability of” guilt-by-association” within gene coexpression networks. BMC bioinformatics 6: 1–10. doi: 10.1186/1471-2105-6-227

Wu H, Kirita Y, Donnelly EL, Humphreys BD. 2019. Advantages of single-nucleus over single-cell RNA sequencing of adult kidney: rare cell types and novel cell states revealed in fibrosis. Journal of the American Society of Nephrology 30: 23–32. doi: 10.1681/ASN.2018090912

Wu T, Hu E, Xu S, Chen M, Guo P, Dai Z, Feng T, Zhou L, Tang W, Zhan L. 2021. clusterProfiler 4.0: A universal enrichment tool for interpreting omics data. The Innovation 2: 100141. doi: 10.1016/j.xinn.2021.100141

Yu H, Luscombe NM, Qian J, Gerstein M. 2003. Genomic analysis of gene expression relationships in transcriptional regulatory networks. Trends in Genetics 19: 422–427. doi:10.1016/S0168-9525(03)00175-6

Zhang L, Li W-H. 2004. Mammalian housekeeping genes evolve more slowly than tissue-specific genes. Molecular biology and evolution 21: 236–239. doi: 10.1093/molbev/msh010

Zhu J, He F, Hu S, Yu J. 2008. On the nature of human housekeeping genes. Trends in genetics 24: 481–484. doi: 10.1016/j.tig.2008.08.004

