## Supplementary Figures for "In search of a core cellular network with single cell transcriptome data in the fruit fly"

Figure S1

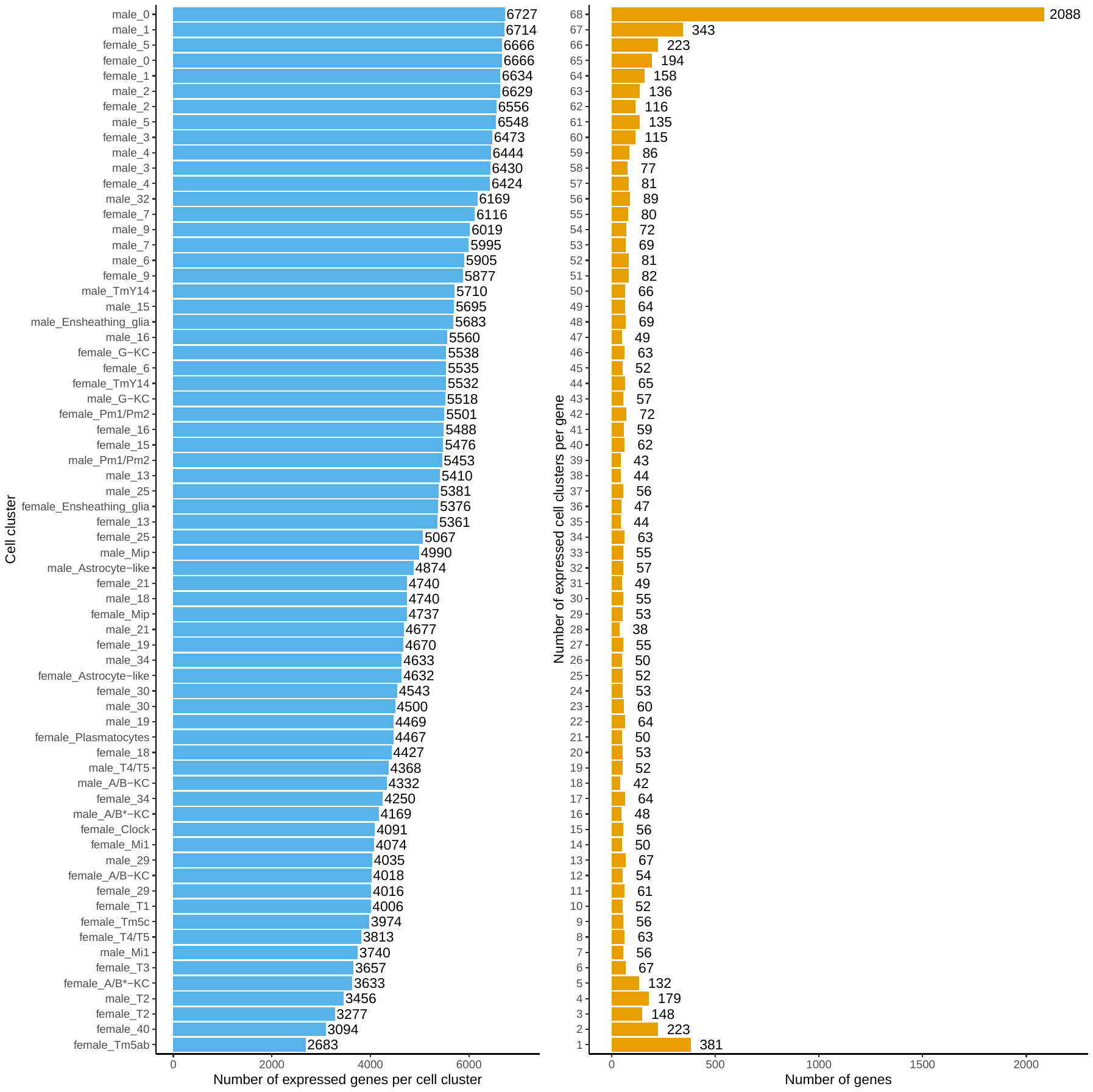

**Figure S1. Number of expressed genes per cell cluster and the number of expressed cell clusters per gene.**

The number of expressed genes for each brain cell cluster (left) and the number of cell clusters one gene was detected as expressed (right).

Figure S2

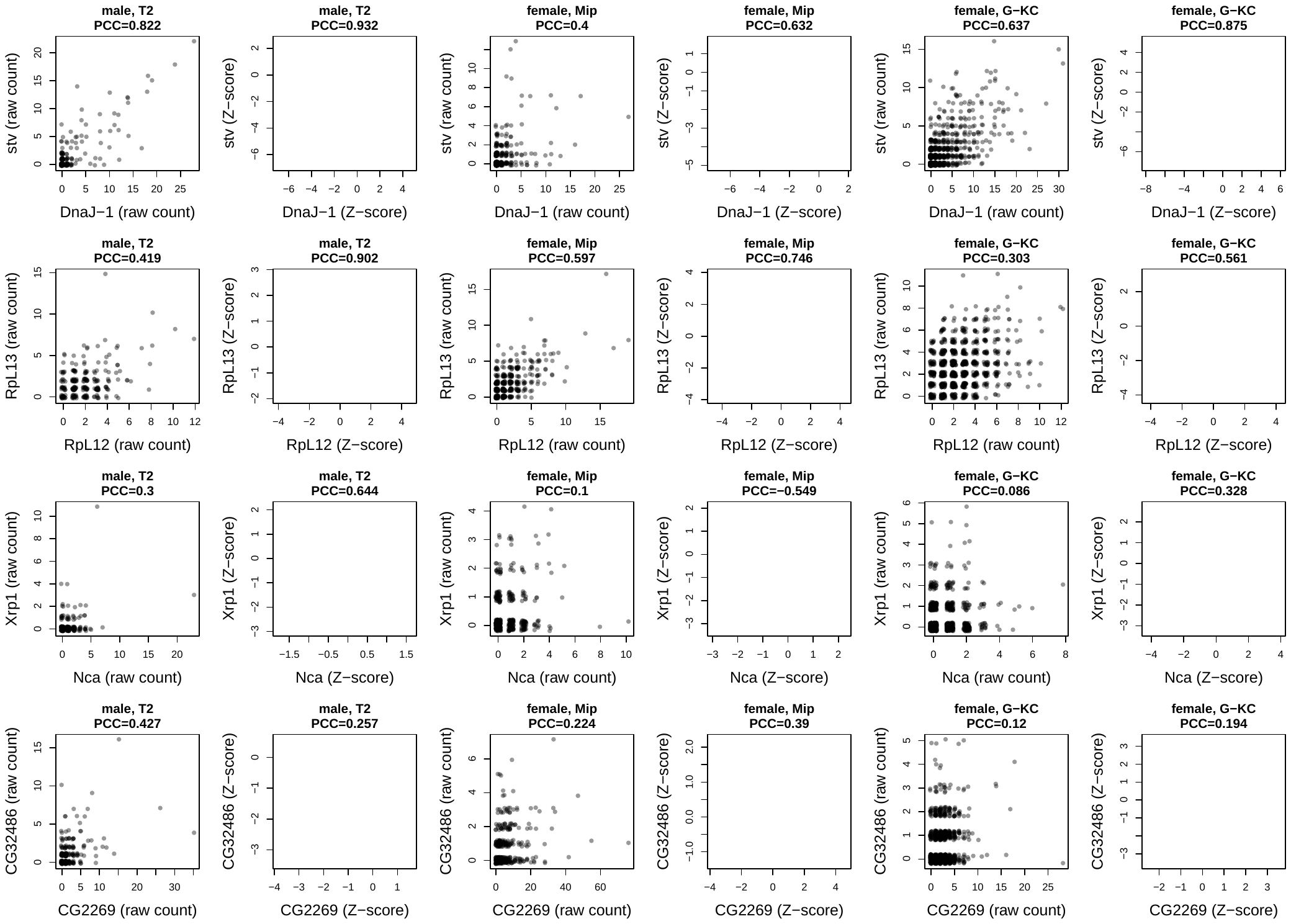

**Figure S2. Example scatter plots of four gene pairs in three cell clusters**.

The bigScale2 algorithm transforms raw gene count values into Z-scores before measuring gene associations, this figure shows the scatter plots of four gene pairs using raw count data or Z-score transformed values. Each row corresponds to one gene pair, each column is one cell cluster. Black dots indicate raw count data and blue dots Z-score transformed data. The cell cluster annotation and the Pearson Correlation Coefficient value for the respective scatter plot are shown on top of each panel figure.

Figure S3

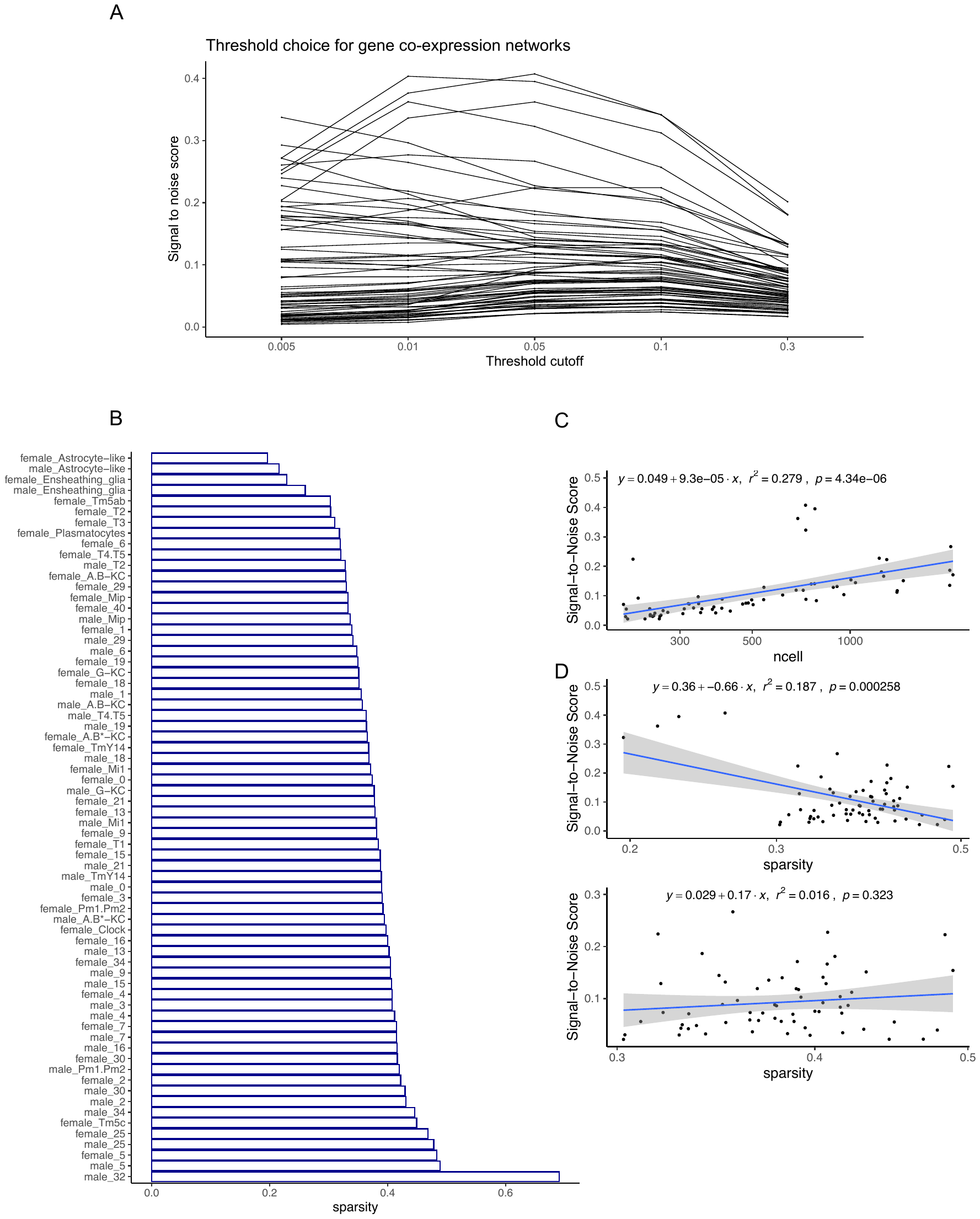

**Figure S3. Use signal-to-noise scores to select a thresholding value for gene correlation matrices and to assess the effect of number of cells and sparsity on network consistency.**

A. The signal to noise score at various thresholding values for each cell cluster in the fly brain dataset. The x-axis denotes the top percentile threshold cutoff value used in selecting gene pairs into co-expression networks. Each dot denotes the median signal-to-noise score across 10 subsampling iterations per cell cluster per threshold value. Different lines represent different cell clusters. See **Supplementary Material, section 1** for details.

B. The sparsity levels across the 68 cell clusters in females or males. Sparsity is measured as the percentage of zeros in the gene count matrix at a cell cluster level. The sparsity ranges from 19.63% to 48.91% except cell cluster 32 in males, which has a sparsity level of 69.10%. Cell cluster 32 in males is excluded from further gene co-expression network analysis.

C. The relationship between the number of cells per cluster and its signal-to-noise median score at top 5% thresholding value. A linear regression result is shown and the shading around regression lines indicates 95% confidence intervals.

D. The relationship between sparsity per cluster and its signal-to-noise median score at top 5% thresholding value. The upper panel includes all 67 cell clusters and the bottom panel excludes the leftmost 4 data points. A linear regression result is shown in each panel and the shading around regression lines indicates 95% confidence intervals.

Figure S4

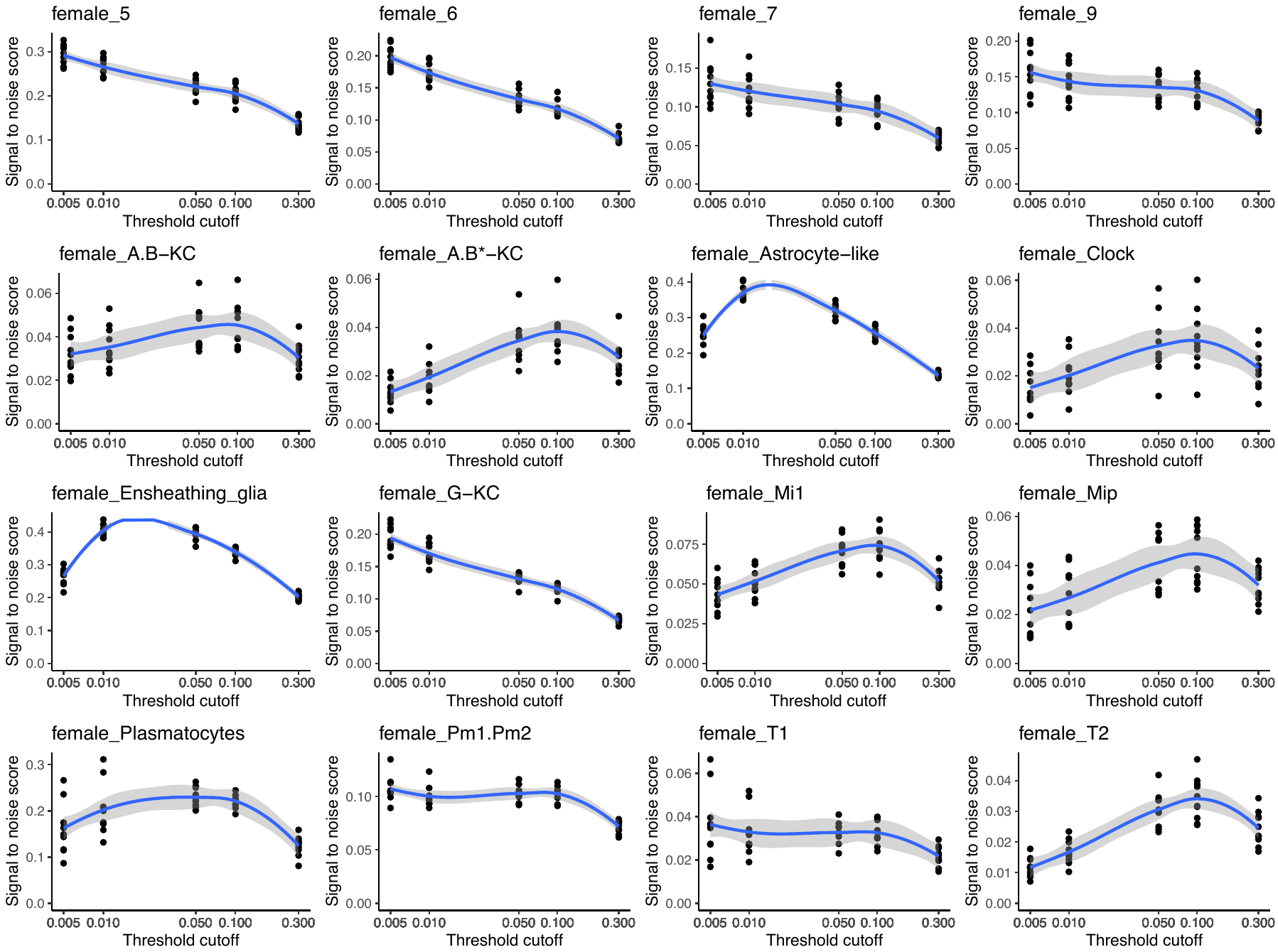

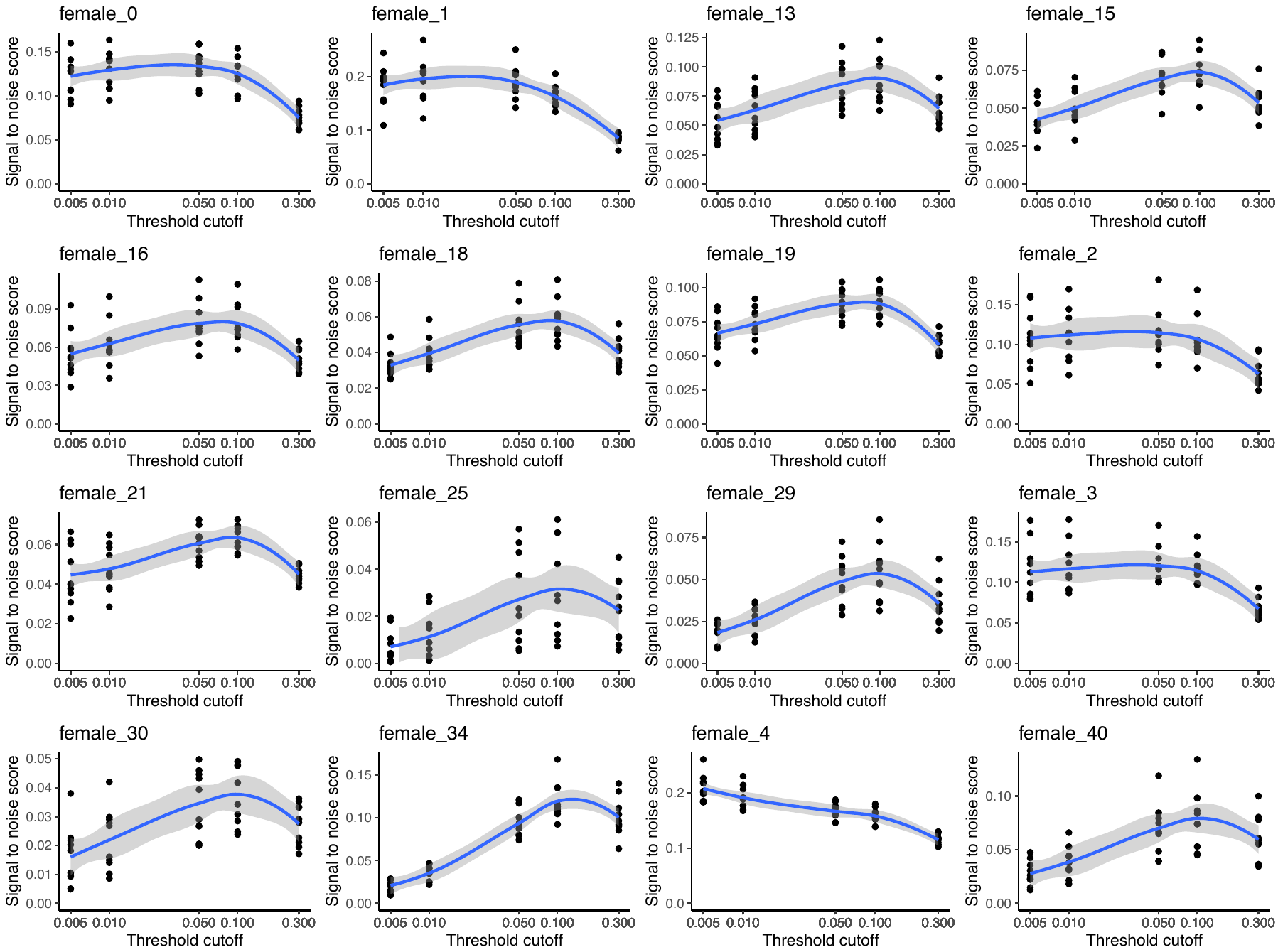

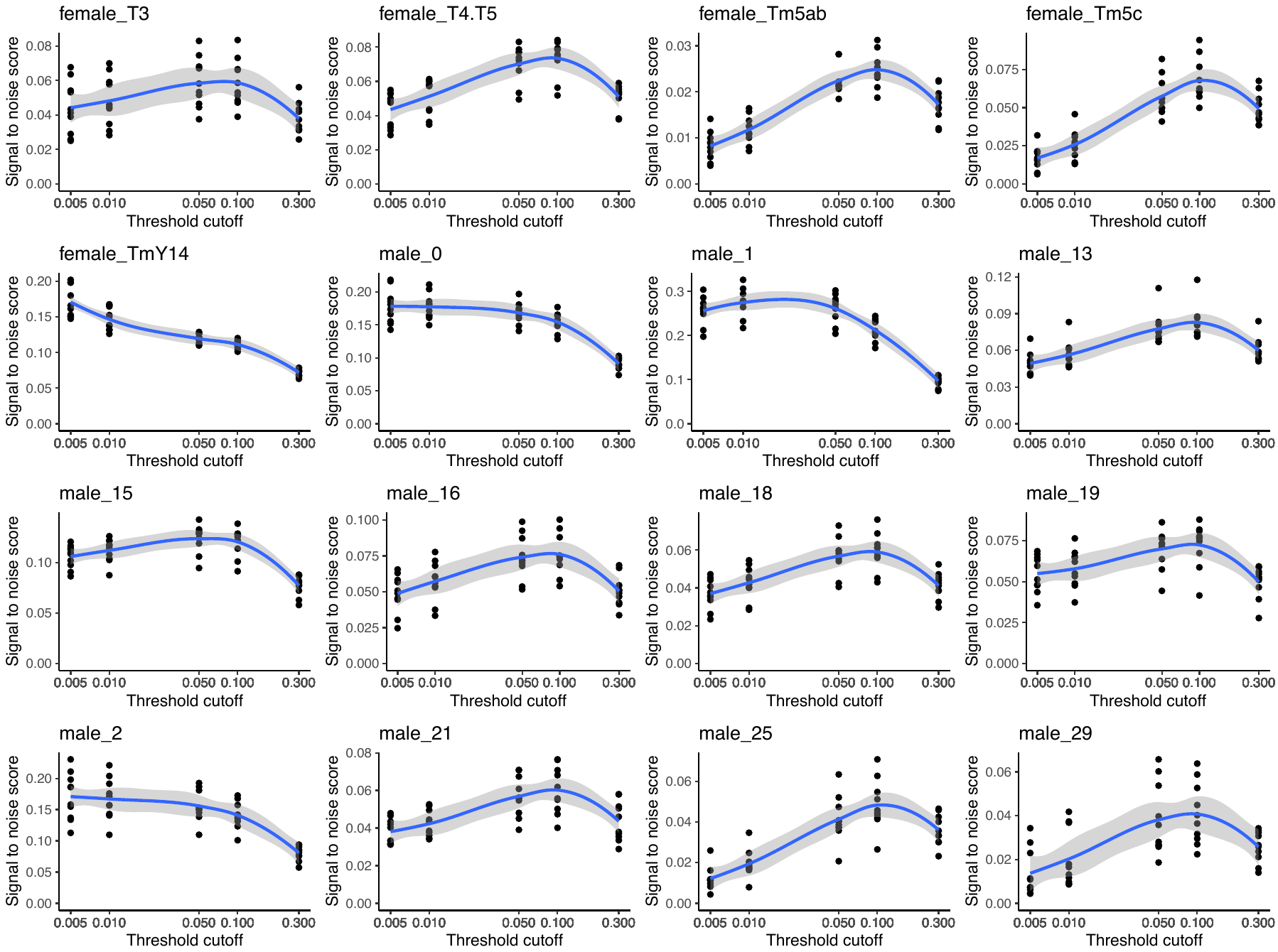

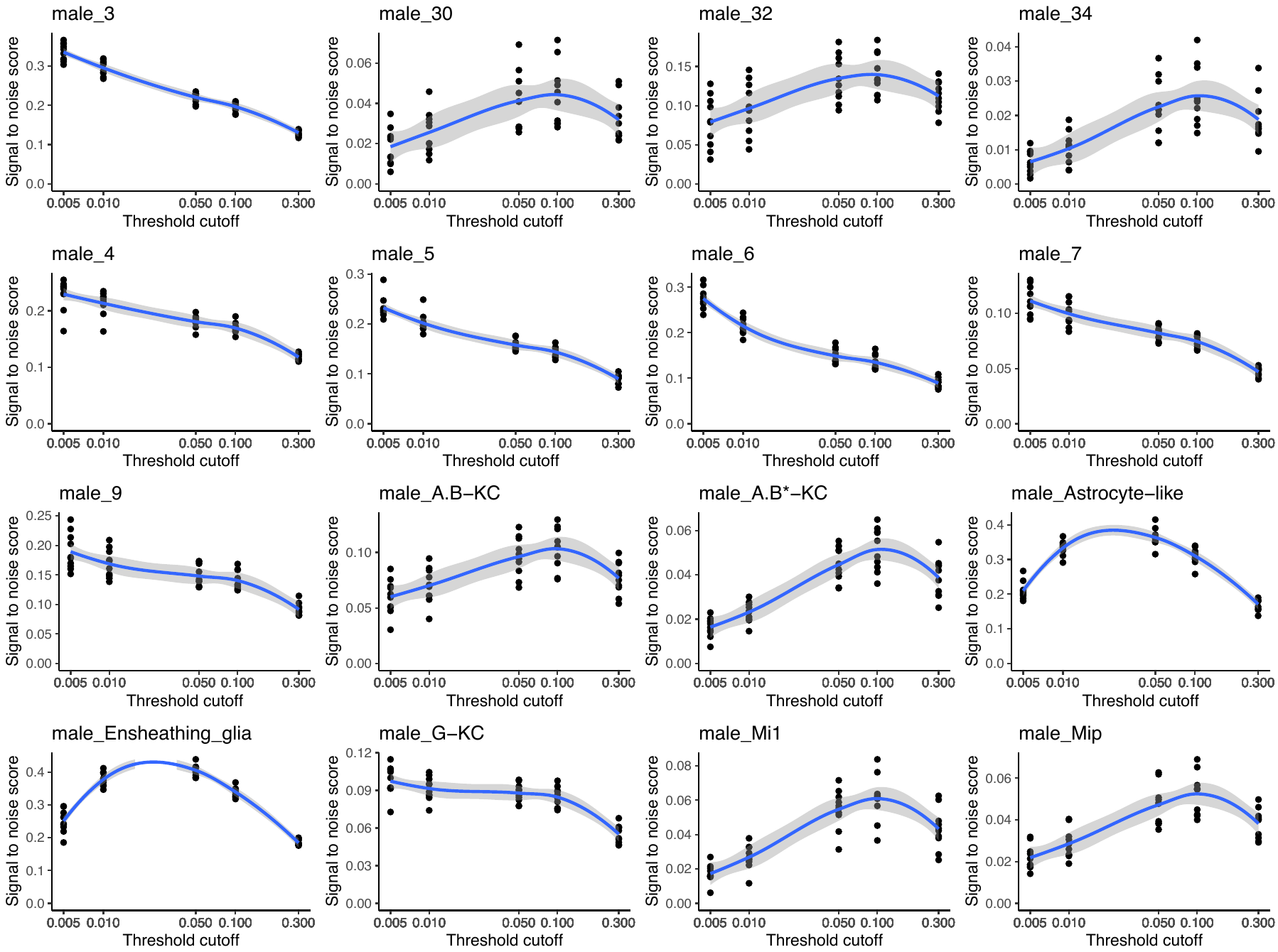

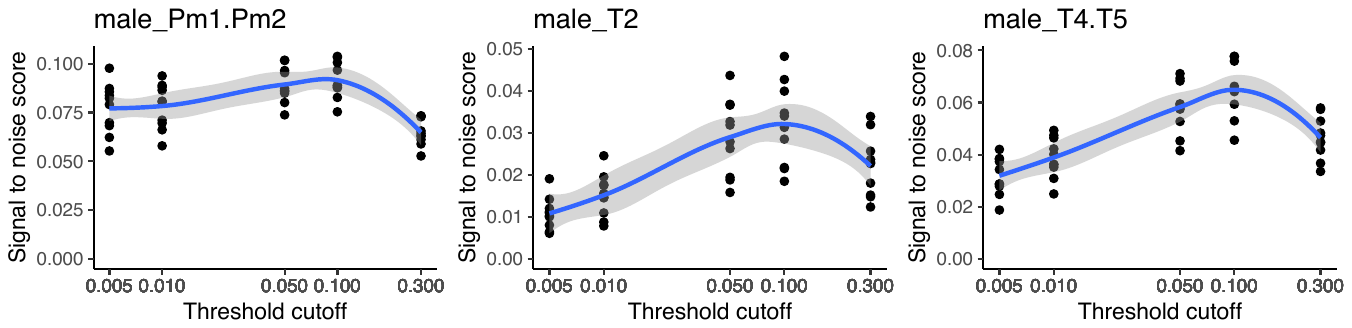

**Figure S4. The signal-to-noise score at various thresholding values for each cell cluster in the fly brain dataset.**

For each panel figure, the x-axis denotes the top percentile threshold cutoff value used in selecting gene pairs into co-expression networks. Each dot denotes a signal-to-noise score from one subsampling iterations per cell cluster per threshold value. The blue line shows the loess regression curve via the geom_smooth function in the R package ggplot2 and the shading around regression lines indicates 95% confidence intervals.

Figure S5

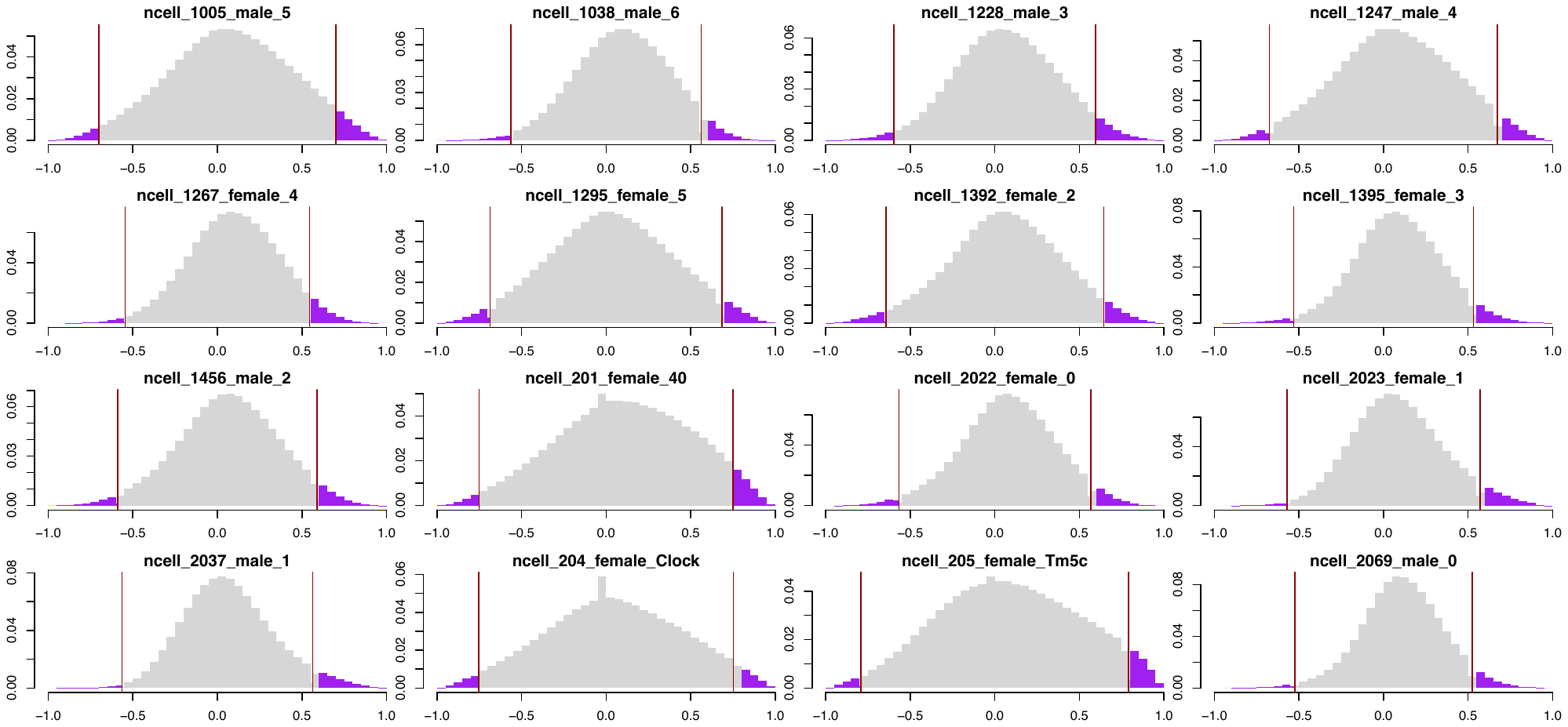

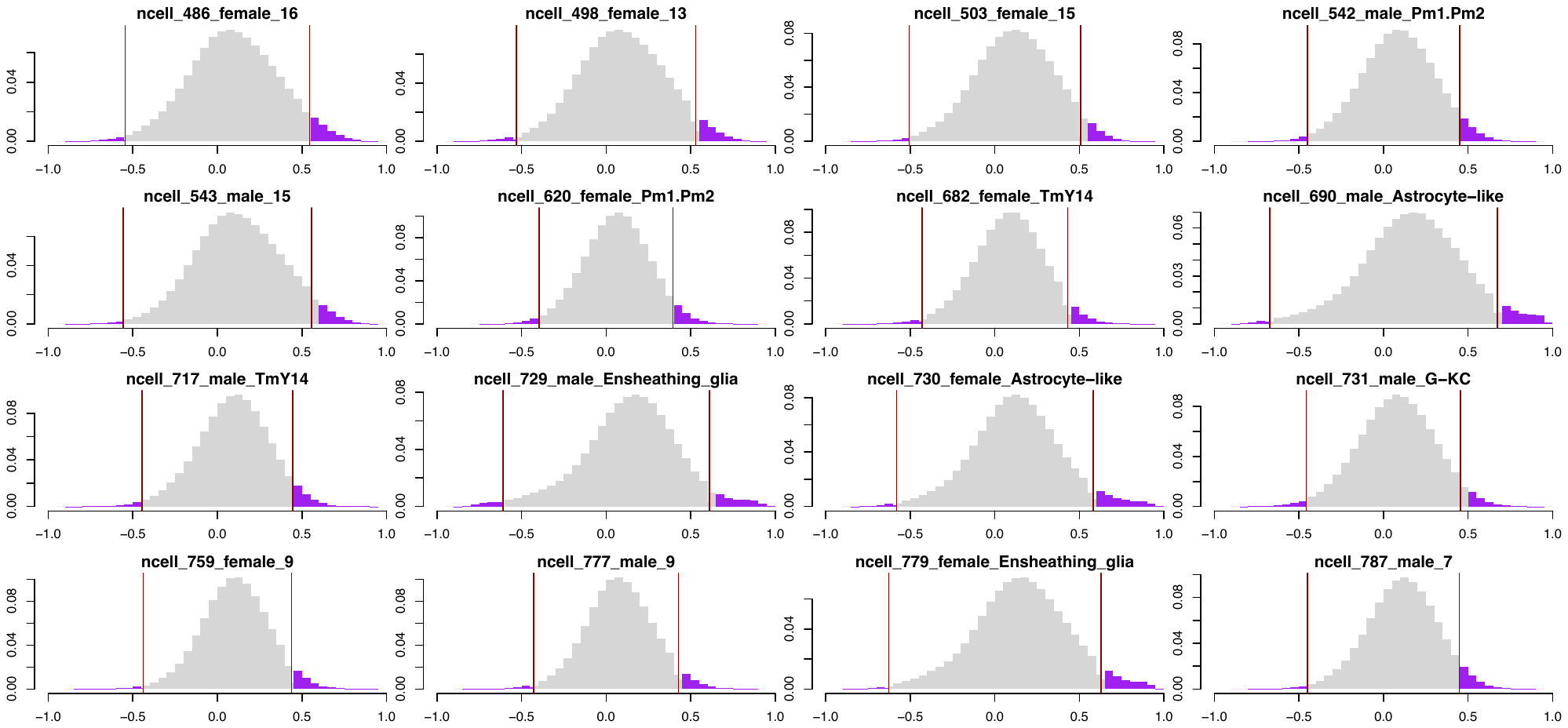

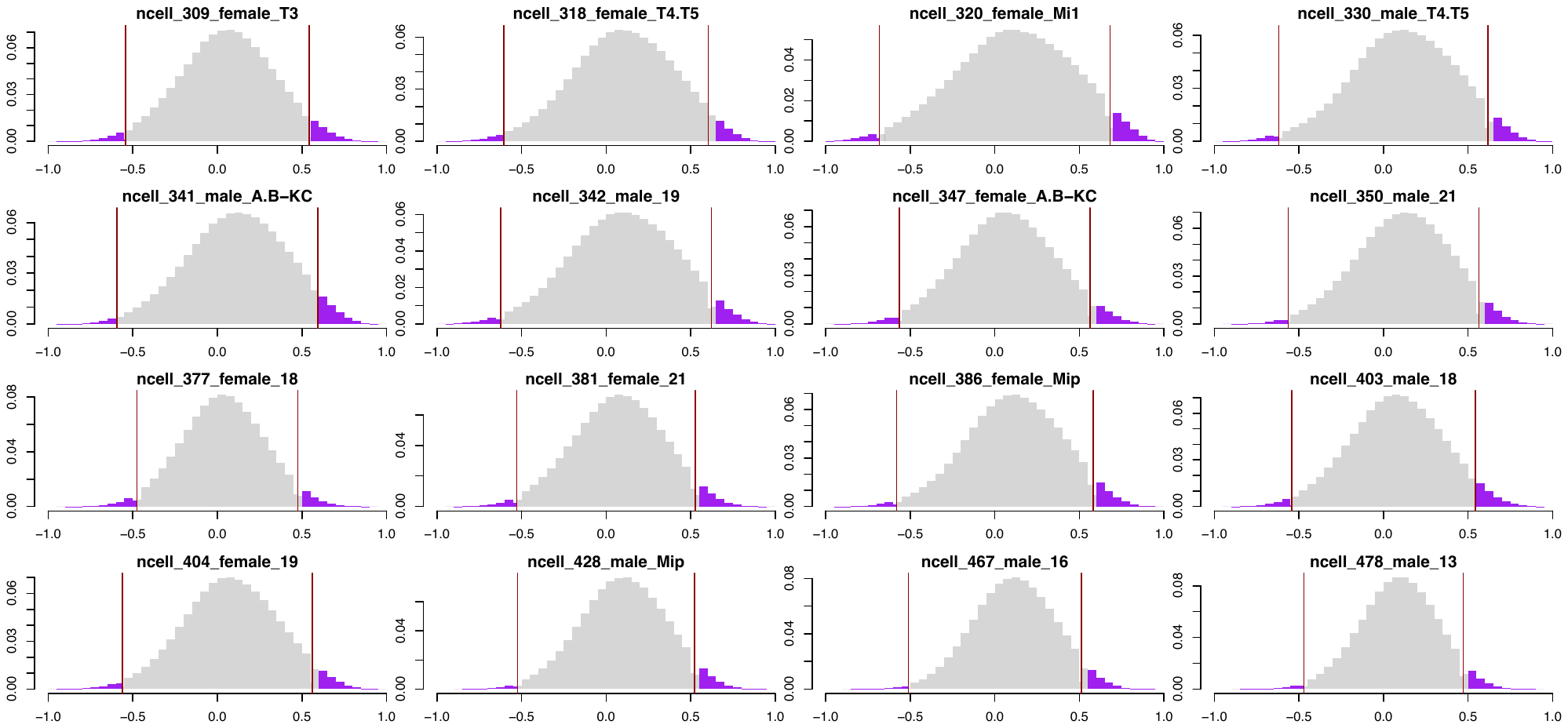

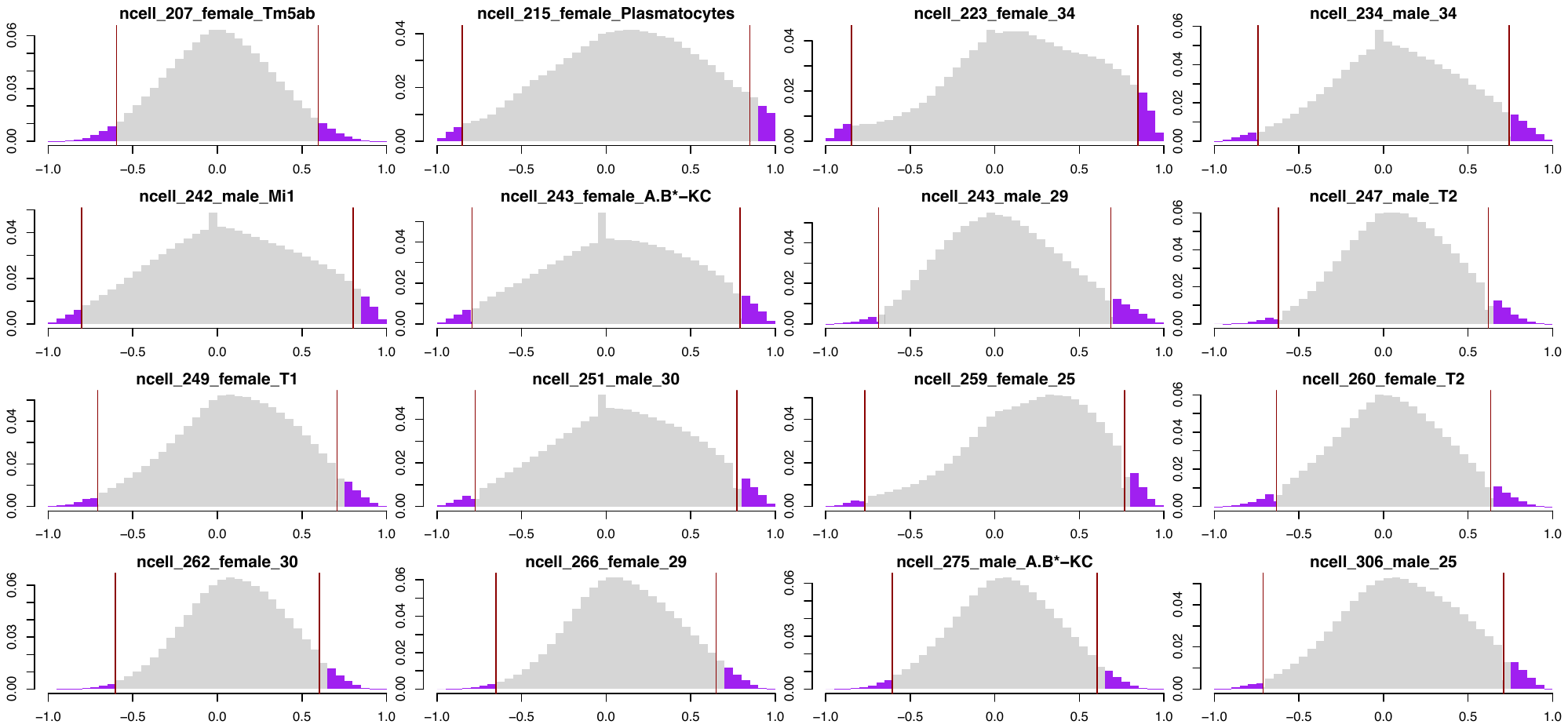

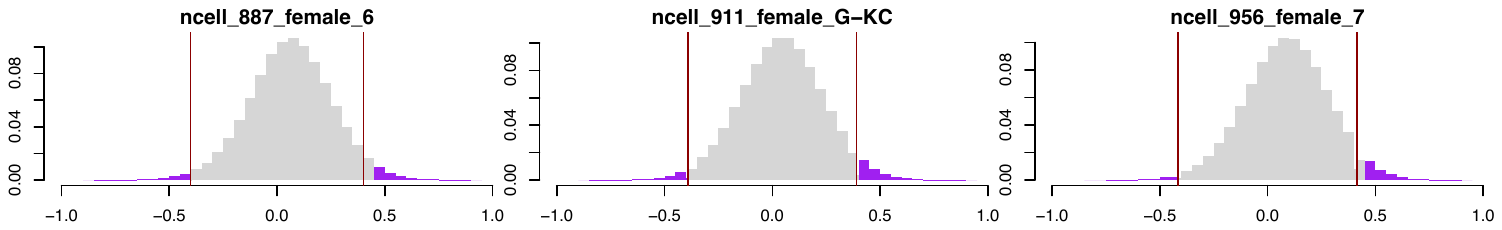

**Figure S5. The Pearson correlation coefficient distribution for each cell cluster.**

The bigScale2 algorithm groups cells into homogeneous clusters and then leverages two genes’ patterns of differential expression between groups to derive the gene-gene correlations using the Pearson correlation coefficient (PCC) method. The resultant PCC distributions followed a bell-curve shape for most cell clusters. The cell cluster annotation is shown on top of each panel figure. The red vertical line indicates the top 5% thresholding values in each cell cluster and the top co-expressed gene pairs are colored in purple.

Figure S6

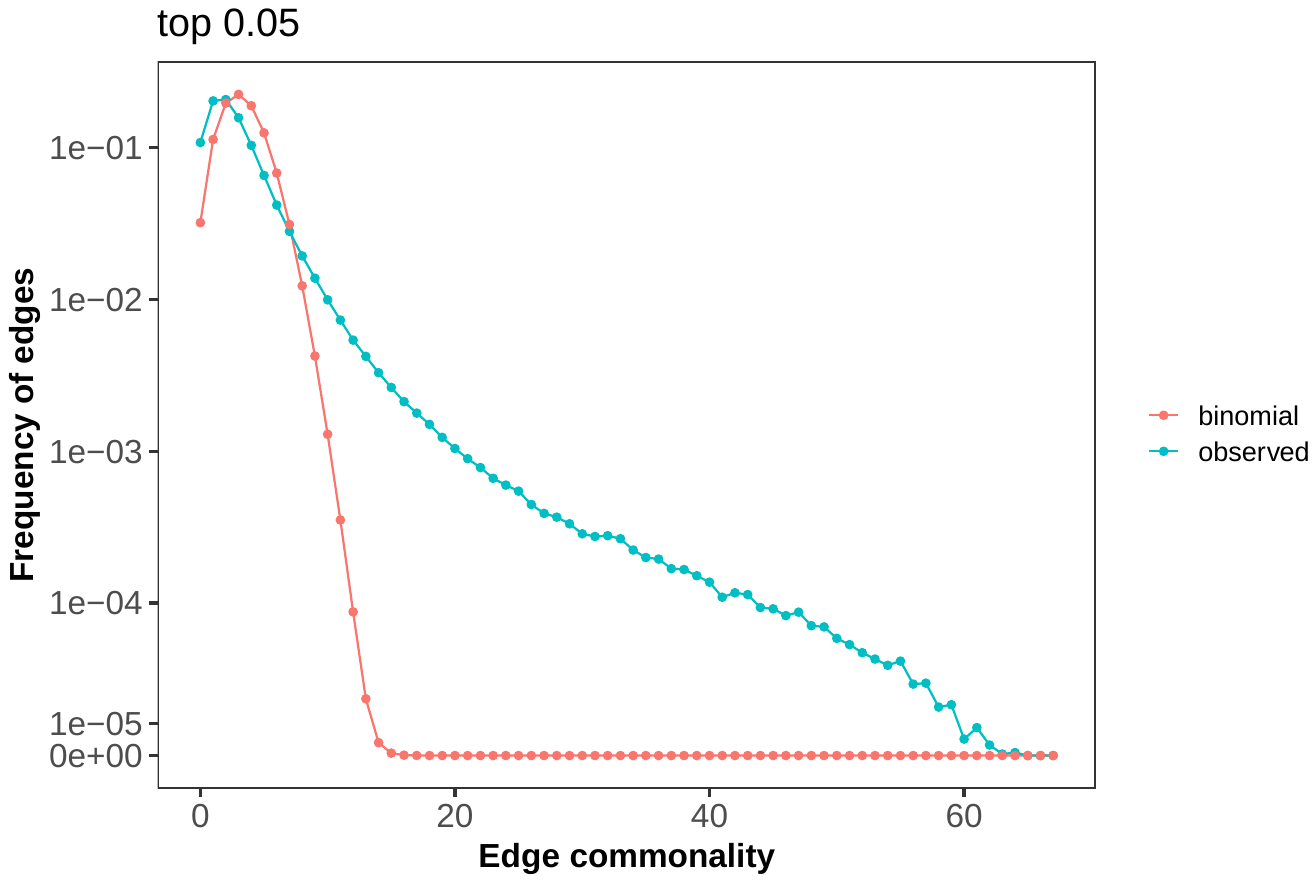

**Figure S6. Comparing the observed edge commonality distribution with the binomial distribution derived approximation.**

The blue dots show the observed edge commonality distribution and the red dots the null distribution from the binomial distribution derived approximation.

Figure S7
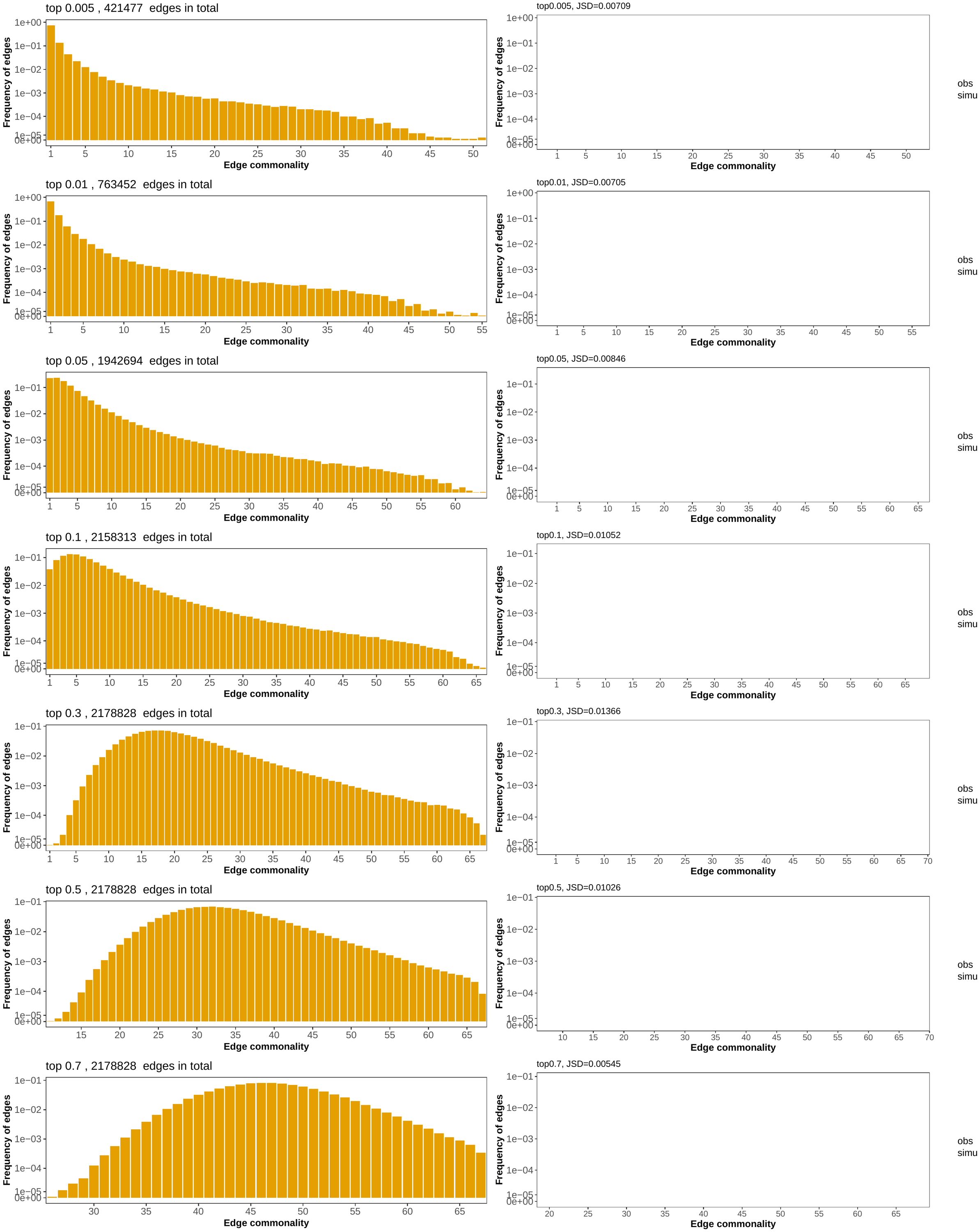

**Figure S7. The observed edge commonality distribution compared with the null generated using network randomization with different top percentile cutoffs.**

Different rows represent different top percentile cutoffs. For each row, the left panel shows the observed edge commonality distribution and the right panel shows the observed edge commonality distribution compared with the null generated using network randomization. For the right panels, the yellow dots show the observed edge commonality distribution and the gray dots the null distribution from network randomization. From top to bottom, the percentile cutoff values used in network construction change from stringent to loose: 0.005, 0.01, 0.05, 0.1, 0.3, 0.5, and 0.7. Network randomization was performed 20 times for each cell cluster individually with network size (number of nodes and edges) and gene degree (number of co-expressed gene partners per gene) fixed. The distance between the observed and null distributions was measured using the Jensen–Shannon divergence (JSD) metric and shown on top of each panel figure.

Figure S8

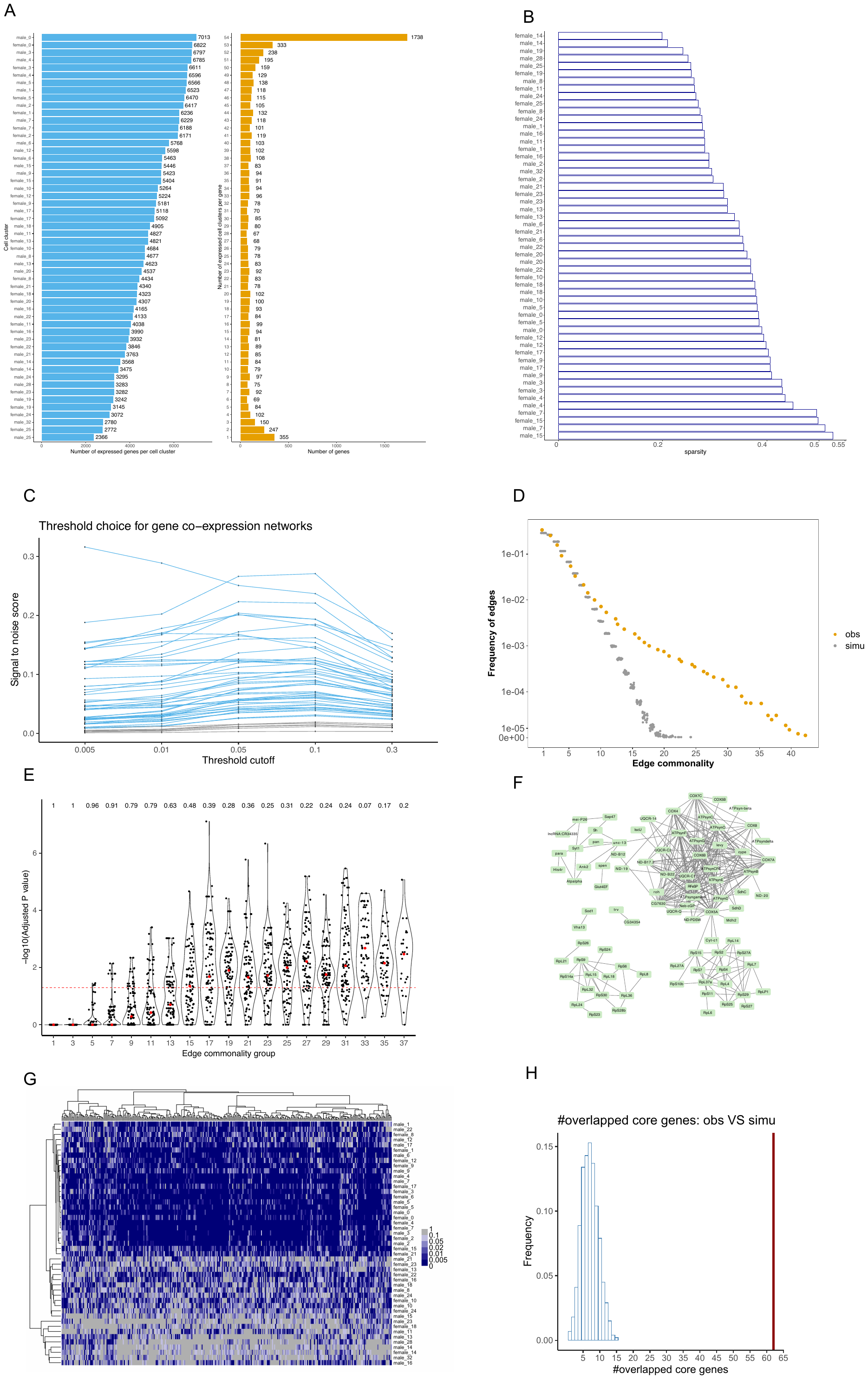

**Figure S8. Analysis of the single-cell dataset of fly brains** **from** **Baker et al. (2021)**

A. Number of expressed genes per cell cluster and the number of expressed cell clusters per gene. The number of expressed genes for each brain cell cluster (left) and the number of cell clusters one gene was detected as expressed (right). See **Supplementary Material, section 2** for details.

B. The sparsity levels across cell clusters in females or males.

C. The signal to noise score at various thresholding values for each cell cluster. The x-axis denotes the top percentile threshold cutoff value used in selecting gene pairs into co-expression networks. Each dot denotes the median signal-to-noise score across 10 subsampling iterations per cell cluster per threshold value. Different lines represent different cell clusters. Blue lines represent cell clusters with a minimal signal to noise score 0.02 which is utilized in following gene co-expression network analysis.

D. The observed edge commonality distribution (yellow) compared with the null expectation derived from network randomization (gray). Network randomization was performed 20 times for each cell cluster individually with network size (number of nodes and edges) and gene degree (number of co-expressed gene partners per gene) fixed.

E. Adjusted P-value distributions of sampled edges in different edge commonality groups using a rank aggregation method. Edge commonality groups were selected to cover the full range of edge commonality scores. Each violin plot illustrates the distribution of adjusted P values for the sampled edges for each edge commonality group. Each dot represents an edge. The dashed red line indicates an adjusted P value of 0.05. The number above each violin indicates the percentage of edges with adjusted P values smaller than 0.05 in the corresponding edge commonality group.

F. Network visualization of the core network.

G. Heatmap of the normalized ranks for the 323 edges with a commonality score at least 33 across cell clusters. Each edge has a rank *r* in each cell cluster based on its absolute correlation value, from which a normalized rank r’ = (r-1)/(R-1) is calculated, where R is the total number of edges in a cell network. A smaller normalized rank value indicates this edge is highly ranked among gene pairs and is encoded as dark blue colors in the heatmap. The row labels are cell cluster annotations and each column is a gene pair. The trees on both sides are generated via hierarchical clustering using the ‘hclust’ function in R with the "ward.D2" method on the Euclidean-based distances.

H. The number of overlapped genes between core networks from single-cell fly brain dataset in the main text and from Baker et al. (2021) is significant compared with simulations. There were 2088 commonly expressed genes in single-cell and 1738 in the Baker et al. (2021) dataset, respectively. The number of core genes in the single-cell dataset is 205 and in the Baker et al. (2021) dataset 88, between which 62 genes overlapped. To assess its significance, we performed simulations. In each simulation replicate, 205 genes were randomly sampled from the 2088 commonly expressed genes and 88 genes from the 1738 genes, respectively, and the number of overlapped genes was recorded. The figure shows the distribution of the number of overlapped genes from 1000 simulations. The red vertical line indicated the number of observed overlapped genes.

Figure S9

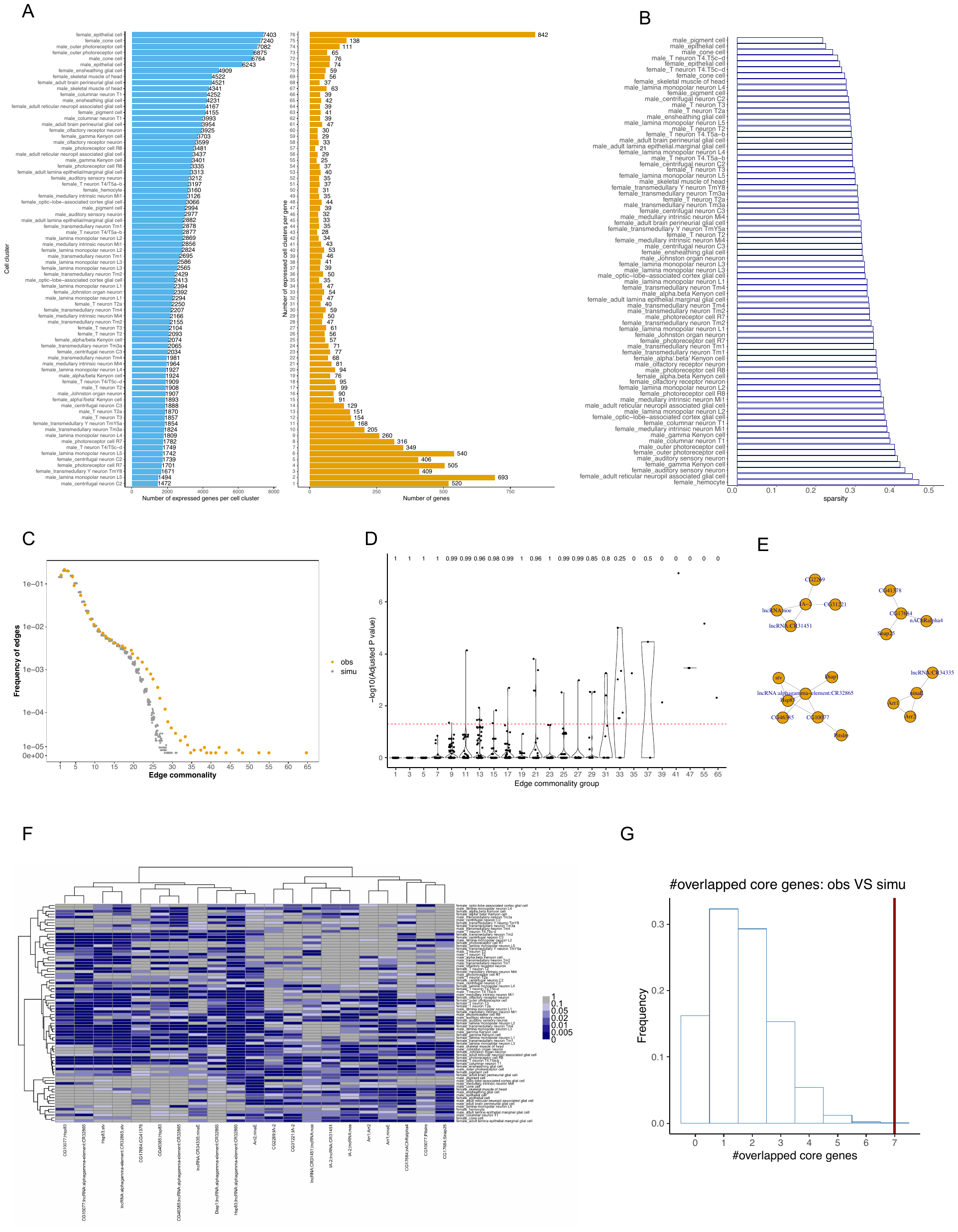

**Figure S9. Analysis of the single-nucleus dataset from fly heads from Li et al. (2021)**

A. Number of expressed genes per cell type and the number of expressed cell types per gene. The number of expressed genes for each brain cell type (left) and the number of cell types one gene was detected as expressed (right). See **Supplementary Material, section 3** for details.

B. The sparsity levels across the 76 cell types in females or males.

C. The observed edge commonality distribution (yellow) compared with the null expectation derived from network randomization (gray). Network randomization was performed 100 times for each cell type individually with network size (number of nodes and edges) and gene degree (number of co-expressed gene partners per gene) fixed.

D. Adjusted P-value distributions of sampled edges in different edge commonality groups using a rank aggregation method. Edge commonality groups were selected to cover the full range of edge commonality scores. Each violin plot illustrates the distribution of adjusted P values for the sampled edges for each edge commonality group. Each dot represents an edge. The dashed red line indicates an adjusted P value of 0.05. The number above each violin indicates the percentage of edges with adjusted P values smaller than 0.05 in the corresponding edge commonality group.

E. Network visualization of the core network.

F. Heatmap of the normalized ranks for the 21 edges with a commonality score at least 35 across cell types. Each edge has a rank *r* in each cell type based on its absolute correlation value, from which a normalized rank r’ = (r-1)/(R-1) is calculated, where R is the total number of edges in a cell network. A smaller normalized rank value indicates this edge is highly ranked among gene pairs and is encoded as dark blue colors in the heatmap. The row labels are cell type annotations and the column labels are gene pairs. The trees on both sides are generated via hierarchical clustering using the ‘hclust’ function in R with the "ward.D2" method on the Euclidean-based distances.

G. The number of overlapped genes between core networks from single-cell fly brain dataset in the main text and from single-nucleus fly head dataset is significant compared with simulations. There were 2088 commonly expressed genes in single-cell and 842 in single-nucleus datasets, respectively. The number of core genes in the single-cell dataset is 205 and in the single-nucleus dataset 20, between which 7 genes overlapped. To assess its significance, we performed simulations. In each simulation replicate, 205 genes were randomly sampled from the 2088 commonly expressed genes and 20 genes from the 842 genes, respectively, and the number of overlapped genes was recorded. The figure shows the distribution of the number of overlapped genes from 1000 simulations. The red vertical line indicated the number of observed overlapped genes.

Figure S10

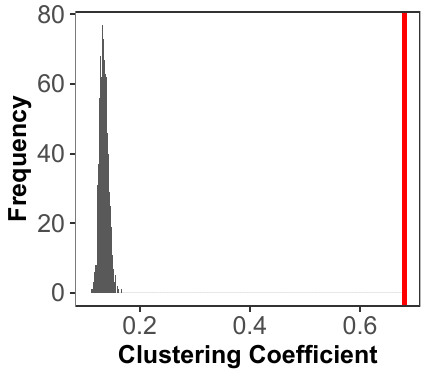

**Figure S10. Clustering coefficient of the core network compared with the null distribution.**

The core network has a clustering coefficient of 0.68, indicated by the red vertical line. To generate a null distribution of the clustering coefficient, we randomly sampled an equal number of genes and edges as the observed core network while keeping the gene degree distribution fixed. We randomly sampled 1000 times and calculated each corresponding clustering coefficient value. The null distribution shows that the maximal clustering coefficient of the null is 0.16 while the observed one is 0.68.

Figure S11

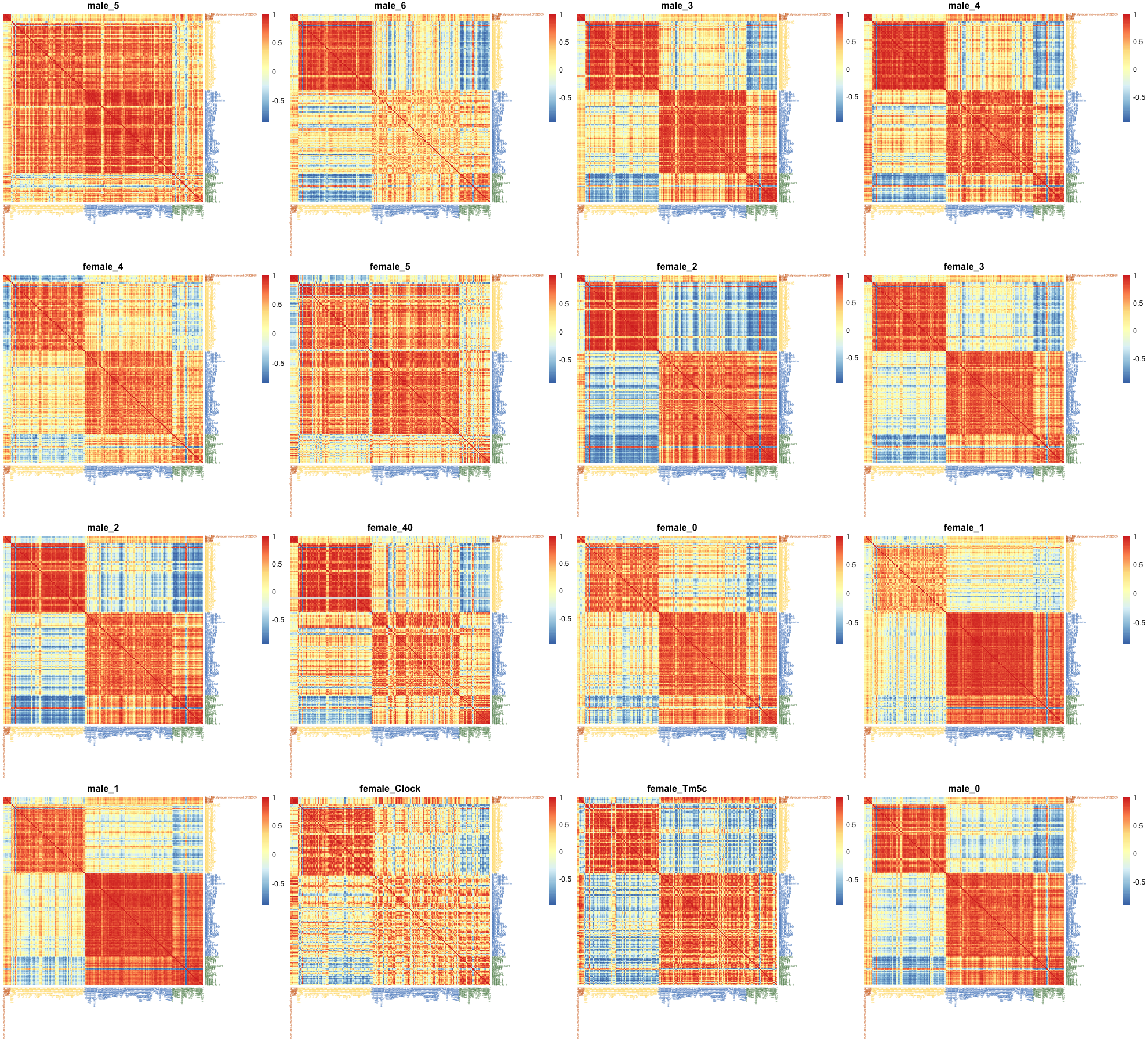

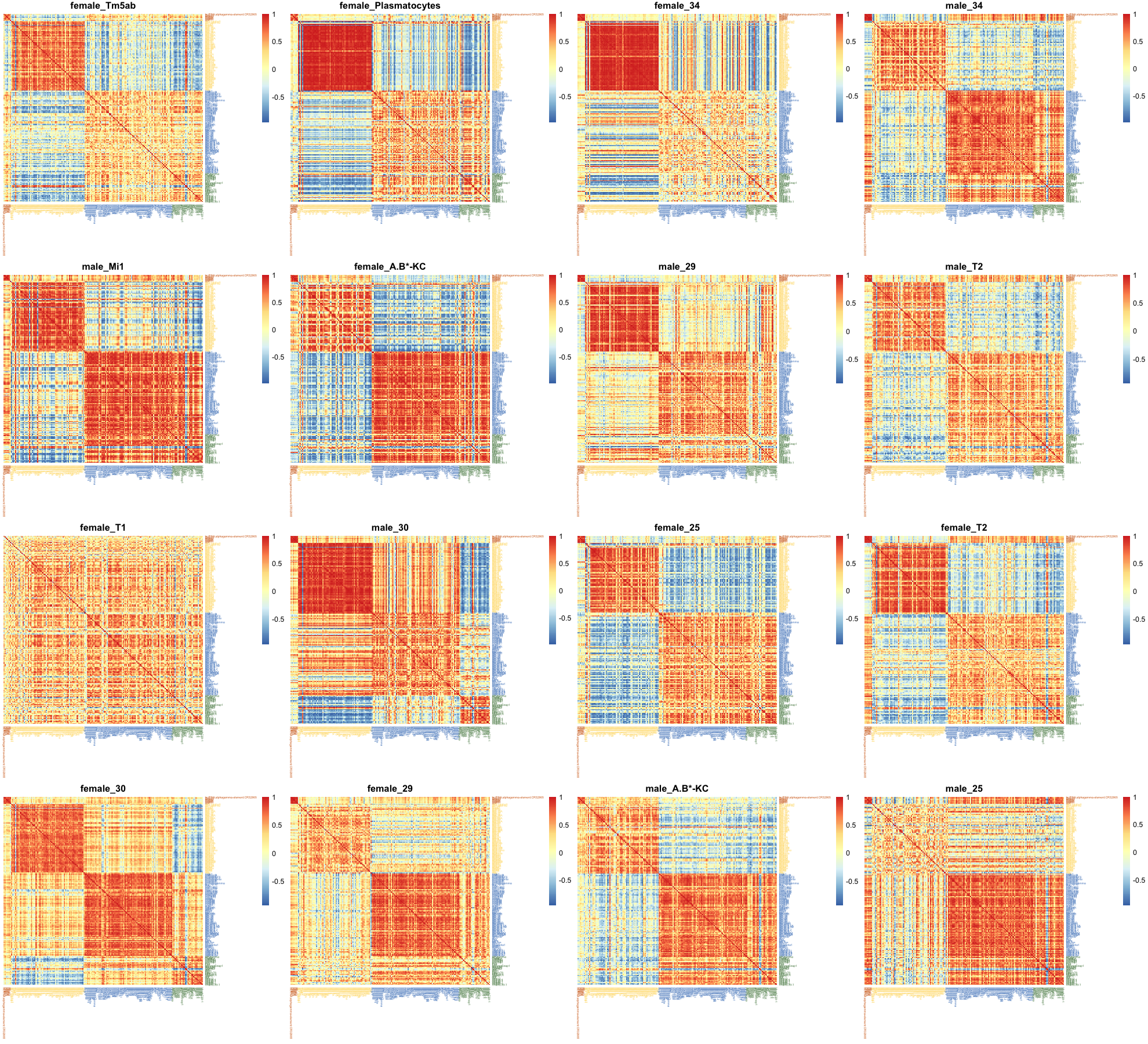

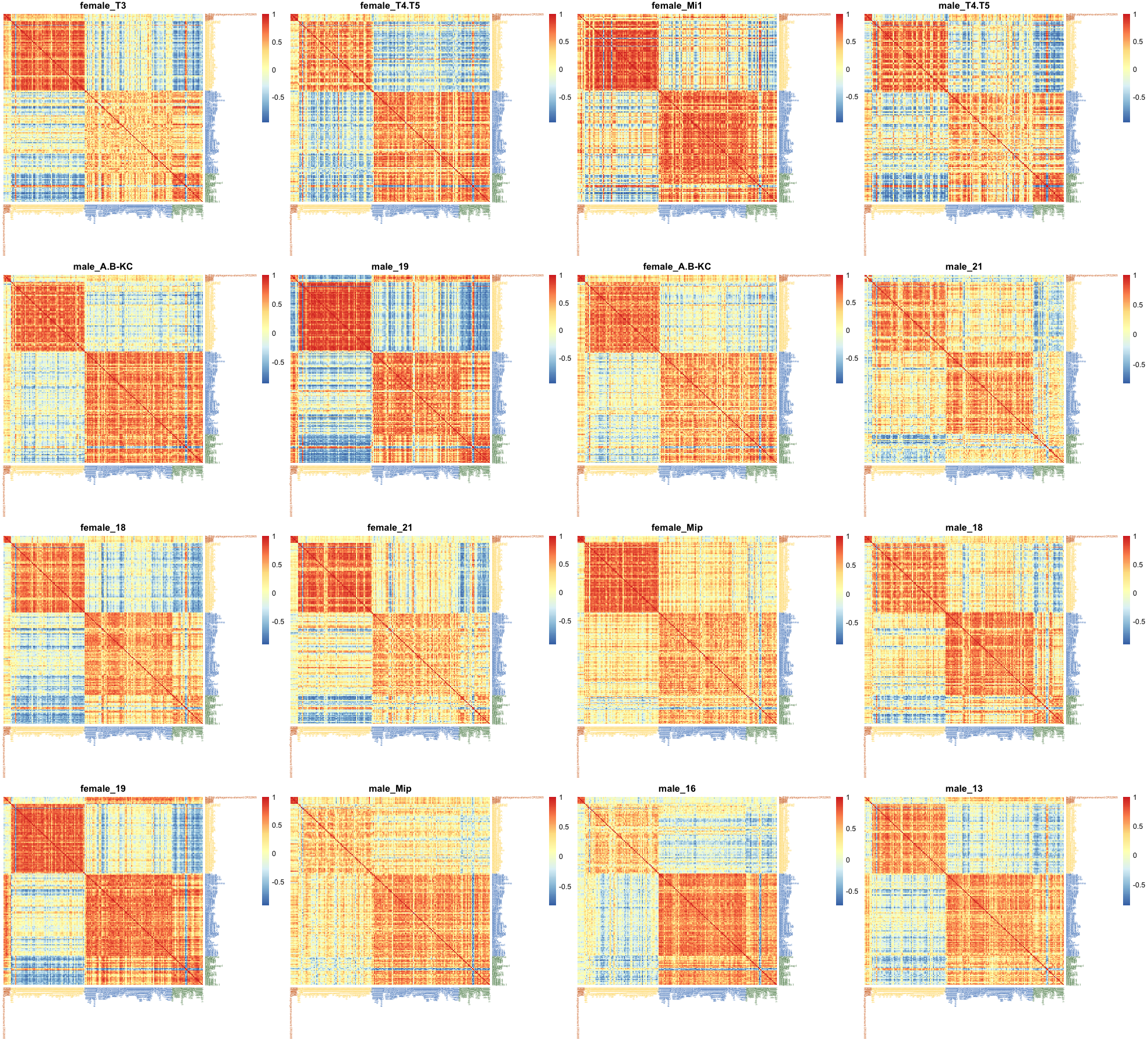

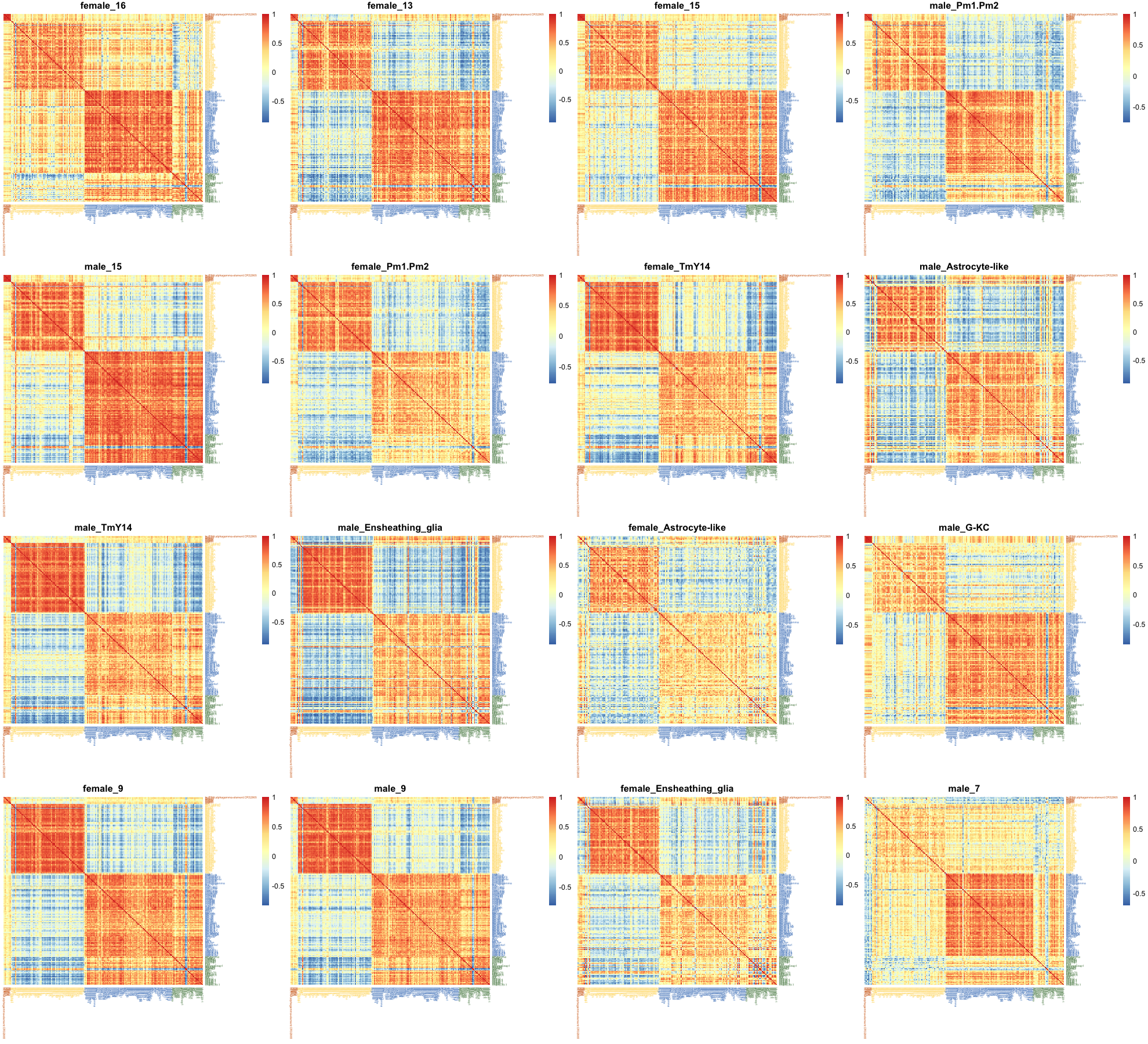

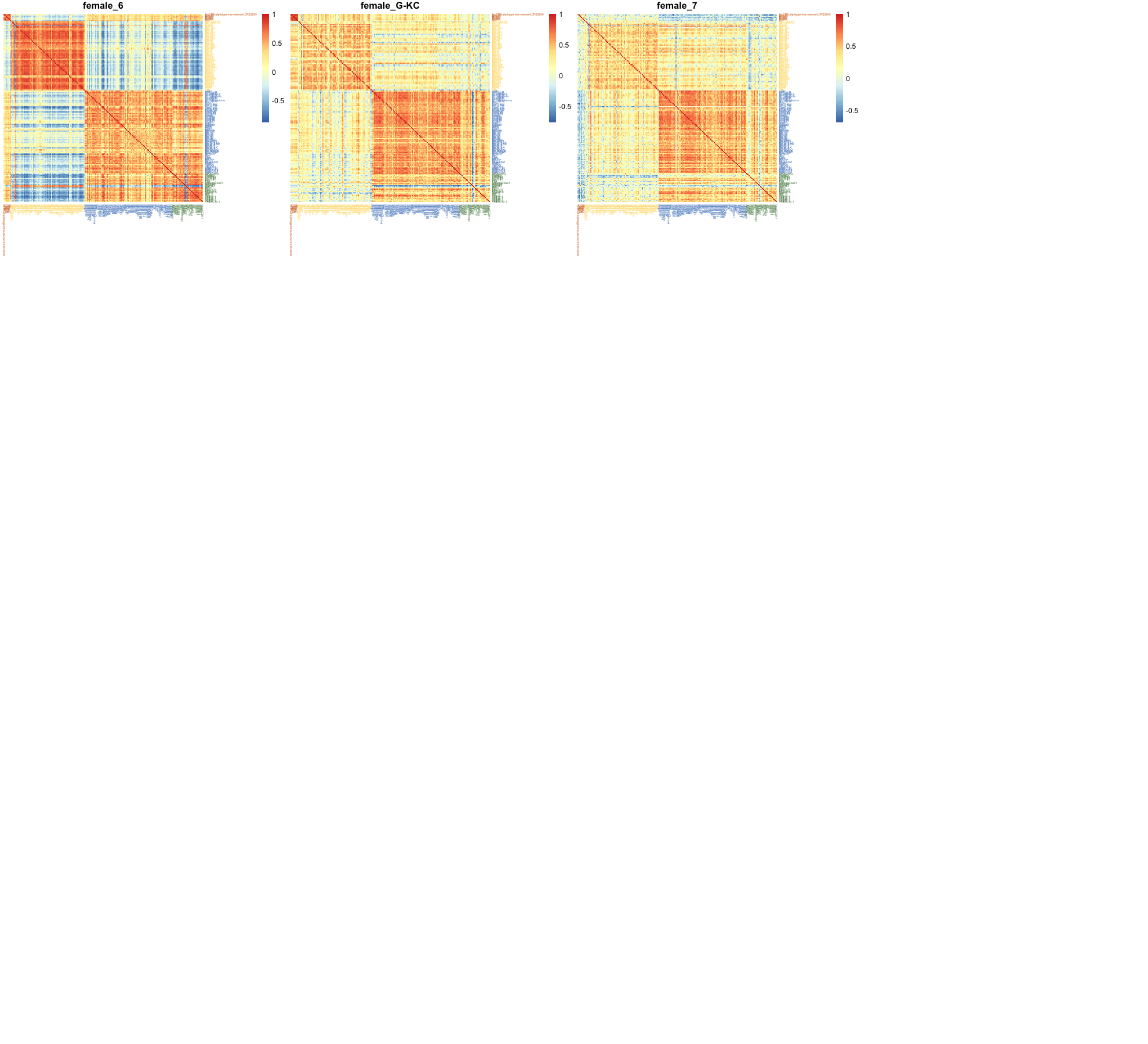

**Figure S11. Heatmap of gene correlations of core modules in each cell cluster.**

Pairwise gene correlations of core module gene members are shown for each cell cluster separately. Gene names are shown in the row and column labels whose colors correspond to core modules described in the main text.

Figure S12
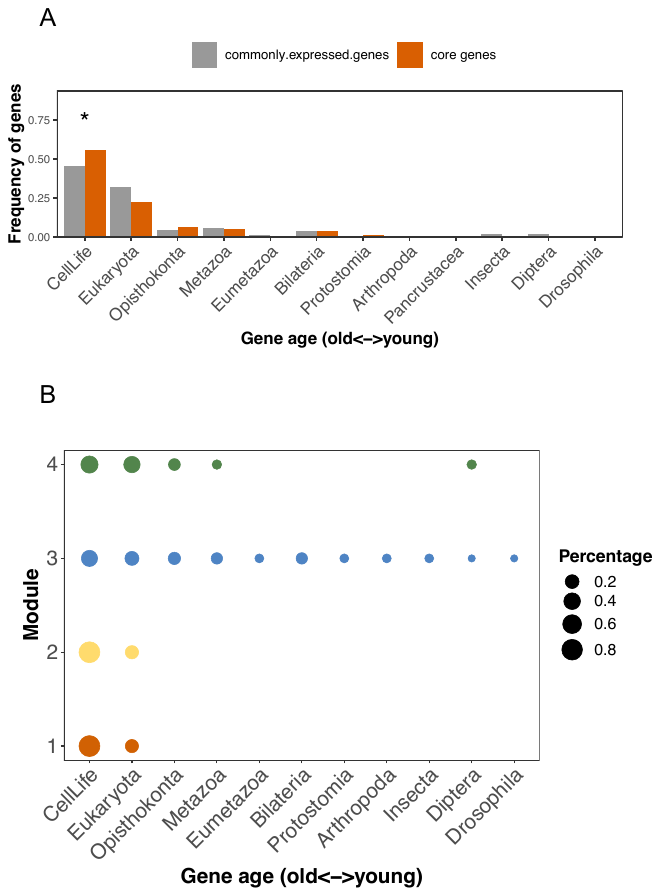

**Figure S12. Evolutionary signatures of core modules.**

A. We assigned fly genes into different evolutionary age groups in a phylostratigraphy framework. The number of genes in each evolutionary age group were compared between commonly expressed genes and genes constituting the core using a one-sided fisher exact test. Multiple-testing was adjusted using the p.adjust function in R with the Benjamini & Hochberg method. *, p.adjust < 0.05.

B. The age distribution of gene members of each core module. The size of each circle represents the proportion of genes in that evolutionary age group in the corresponding module.
