## Supplementary Materials for "In search of a core cellular network with single cell transcriptome data in the fruit fly"

**Supplementary Material**

**Table of contents:**

Section 1. Selecting a thresholding value in correlation matrix and evaluate the performance of the bigScale2 algorithm for data sparsity

Section 2. Analysis of cell clusters in the fly brain single-cell RNA-seq dataset from Baker et al. (2021)

Section 3. Analysis of cell types in the fly head single-nucleus RNA-seq dataset from Li et al. (2021)

**Section 1. Selecting a thresholding value in correlation matrix and evaluate the performance of the bigScale2 algorithm for data sparsity**

*Selecting a thresholding value in correlation matrix*

In thresholding a correlation matrix, we want to find a threshold value that minimizes the number of edges between genes that are not co-expressed, and maximizes the number of edges between genes that are co-expressed. While no consensus exists regarding the choice of the threshold value in gene correlation networks, we tackled this problem using a signal-to-noise score calculated at different threshold values in each cell cluster.

Our procedure starts with the generation of two datasets, through subsampling 75% cells from a given cell cluster. We then constructed two gene co-expression matrices with the bigScale2 algorithm (Iacono et al. 2019). Applying a top percentile threshold cutoff, the two matrices are binarized into two unweighted networks. As these two networks are constructed from a subset of all available cell samples, they are noisier than the network constructed from the full data, however, they should still resemble each other. The similarity of these two networks are measured through a signal-to-noise score approach implemented in the ‘getEdgeSimilarityCorrected’ function from the R package COGENT (Bozhilova et al. 2021). In brief, this function assesses the consistency of two input unweighted networks and calculates a signal-to-noise score by comparing the observed number of overlapped edges to those generated through network randomizations. Specifically, the observed number of overlapped edges is first counted, and then adjusted by the number of overlapped edges from network randomizations. The resultant score ranges from 0 to 1, with a value close to 1 indicating high consistency and to 0 trivial consistency. This procedure is iterated 10 times for every cell cluster at each thresholding value, and each subsampling iteration leads to one signal-to-noise score.

We computed the signal-to-noise score for these 10 iterations for various thresholding values in each cell cluster separately. The median value of the 10 iterations per threshold value per cell cluster was calculated and extracted to be visualized in **Fig. S3A** and individual plot for each cell cluster is shown in **Fig. S4**. The result shows a global trend of signal-to-noise scores changing with thresholding values from different cell clusters, where most cell clusters have the highest score at the top 0.05 threshold cutoff value. We therefore set a thresholding cutoff as the top 5% on gene correlation matrices in our analysis.

*Evaluate the performance of the bigScale2 algorithm for data sparsity*

To evaluate the effect of data sparsity on network construction, we first calculated cell cluster-specific sparsity levels and then examined the relationship between sparsity or other factors and the signal-to-noise scores.

In the fly brain single-cell dataset in the main text, there are 37 cell clusters in females and 31 cell clusters in males which contain at least 200 cells. To identify commonly expressed genes across these cell clusters, we first selected gene sets which were expressed in more than 15 cells and in more than 0.5% of cells for each cell cluster separately, and then extracted their intersections which contained 2088 genes to be referred to as commonly expressed genes. We measured sparsity at a cell cluster level by calculating the percentage of zero in the respective gene count matrix limited to commonly expressed genes. The sparsity ranges from 19.63% to 48.91% except cell cluster 32 in males, which has a sparsity level of 69.10% (**Fig. S3B)**. Cell cluster 32 in males is excluded from further gene co-expression network analysis.

To evaluate the effect of sparsity and other potential factors on network consistency, we extracted the median signal-to-noise scores at the top 5% thresholding cutoff for each cell cluster. Plotting the number of cells against the signal-to-noise scores for each cell cluster shows a significant effect of cell cluster size (P value = 4.34e−06, **Fig. S3C**). Smaller cell clusters tend to have lower signal-to-noise scores while large ones are associated with higher scores.

We went on to examine the effect of sparsity by plotting cell cluster sparsity against their respective signal-to-noise scores. Linear regression using all 67 cell clusters shows a significant relationship (P value = 0.000258, upper panel in **Fig. S3D**). However, this relationship is mainly driven by four cell clusters with the smallest level of sparsity. Indeed, removing these four cell clusters led to a non-significant relationship between sparsity and signal-to-noise score (P value = 0.323, bottom panel in **Fig. S3D**). Thus, the bigScale2 algorithm is not influenced by data sparsity in our dataset.

**Section 2. Analysis of cell clusters in the fly brain single-cell RNA-seq dataset from Baker et al. (2021)**

We downloaded the fly brain atlas data from NCBI Gene Expression Omnibus with GEO accession ‘GSE152495’. This dataset was generated from flies that consumed fixed amounts of sucrose or sucrose supplemented with cocaine, in both sexes, using single-cell libraries on the 10X Genomics platform (Baker et al. 2021). The downloaded dataset contains 8 samples, two replicates per sex per food condition. We integrated the 8 samples following the source code provided by the authors (https://github.com/vshanka23/The-Drosophila-Brain-on-Cocaine-at-Single-Cell-Resolution), and focused our analysis on the 4 samples in female and males under the sucrose food condition. We excluded all mitochondrial genes in the dataset and removed cells that had either less than 200 expressed genes, less than 500 total unique molecular identifier counts, or a total fraction of mitochondrial gene expression exceeding 30%. The filtered dataset contains 10,386 gene expression data in 41,520 high-quality cells grouped into 26 female cell clusters and 28 male cell clusters, with each cell cluster containing at least 200 cells.

To find genes effectively expressed across cell clusters, we selected gene sets which express in more than 15 cells and in more than 0.5% of cells for each cell cluster separately. This procedure led to 1,738 genes identified as commonly expressed genes in these 54 cell clusters (**Fig. S8A**). The sparsity level of all 54 cell clusters were below 53%, measured by the percentage of zeros in the data for the 1,738 commonly expressed genes (**Fig. S8B**).

We used the bigScale2 algorithm (Iacono et al. 2019) to calculate the gene-gene correlation matrix for each cell cluster. We computed signal-to-noise scores at various thresholding values for each cell cluster in order to identify the threshold that maximized this score across most cell clusters. Based on the signal-to-noise score result (**Fig. S8C**), most cell clusters showed the maximal scores at top 5% threshold cutoff. We removed 7 cell clusters with a signal-to-noise score less than 0.02 at the top 5% threshold cutoff, which was the smallest signal-to-noise score at the 5% threshold of the single-cell brain dataset in the main text (**Fig. S3A**). For the remaining 47 cell clusters, we selected the top 5% edges with highest absolute correlation values into the cell cluster-specific networks. Similar to the network analysis of single-cell brain dataset in the main text, the edge commonality distribution shows increasing discrepancy between the observed one and the null generated through network randomization as the edge commonality increases (**Fig. S8D**). This suggests that the real network contains edges that are more common across cells than expected by chance.

We expect to select edges with high commonalities and those which consistently rank highly across the remaining cell clusters where they didn’t make to the top 5% correlations to derive a core network. Similar to the analysis of the single-cell brain dataset in the main text, we used a rank aggregation analysis to inform the edge commonality cutoff. Within each edge commonality group, we randomly sampled 100 edges if this group contained at least 100 edges, otherwise use all available edges. For each edge, we first collected the cell clusters that this edge was absent (didn’t make to the top 5% correlations), and then calculated the adjusted P value on these cell clusters using a rank aggregation method. The result showed that the negative log-transformed adjusted P value distributions increase together with edge commonalities, similar to the pattern observed in the single-cell fly brain atlas dataset, but less pronounced (**Fig. S8E**). In particular, more than 80 percent of edges sampled from the edge commonality groups ≥ 33 were highly ranked in the remaining cell clusters (P < 0.05). Based on this rank aggregation analysis and a P value < 0.05 cutoff, we chose edge commonality 33 as the cutoff to select core network edges, which resulted in a core network in the *Drosophila* head with 88 genes connected by 323 edges (**Fig. S8F)**. We collected all 323 edges and showed their ranks across cell types. Similar to the brain-cell core network in the main text, this core network lacked edges found in all cells, but its edges were highly ranked in the remaining cell clusters (**Fig. S8G**). Compared with the core network which composes 205 genes of the single-cell fly brain dataset in the main text, 62 genes overlapped. Simulations showed significant overlap between core networks inferred from the two datasets (empirical P value = 0.00099, calculated as (n+1)/(N+1), where n indicates the number of permutations that are larger than the observed number and N the total number of permutations, **Fig. S8H)**, validating our discovery in the main text.

**Section 3. Analysis of cell types in the fly head single-nucleus RNA-seq dataset from Li et al. (2021)**

We downloaded the fly head atlas data from the Fly Cell Atlas website (<https://flycellatlas.org/>). This dataset was generated using single-nuclei libraries on the 10X Genomics platform (Li et al. 2021). The downloaded dataset contains 13,056 gene expression data in 100,527 high-quality female or male brain cells grouped into 82 cell clusters. We excluded all mitochondrial genes in the dataset and removed cells that had either less than 200 expressed genes, less than 500 total unique molecular identifier counts, or a total fraction of mitochondrial gene expression exceeding 30%. We excluded two cell clusters labelled as ‘unannotated’ or ‘artefact’, those whose sex labels were ‘mix’, or those with fewer than 200 cells. These procedures led to 40 cell clusters in females and 36 cell clusters in males, all of which were annotated to known head cell types by the authors in the original publication (Li et al. 2021).

To find genes effectively expressed across cell types, we selected gene sets which express in more than 15 cells and in more than 0.5% of cells for each cell type in each sex separately. This procedure led to 842 genes identified as commonly expressed genes in these 76 cell types (**Fig. S9A**). The sparsity level of all 76 cell types were below 50%, measured by the percentage of zeros in the data for the 842 commonly expressed genes (**Fig. S9B**).

The bigScale2 algorithm (Iacono et al. 2019) was employed to calculate the gene-gene correlation matrix and the top 5% edges with highest absolute correlation values were selected to constitute cell type-specific networks. Similar to the analysis of single-cell brain dataset in the main text, the edge commonality distribution shows increasing discrepancy between the observed one and the null generated through network randomization as the edge commonality increases (**Fig. S9C**), suggesting that the real network contains edges that are more common across cells than expected by chance.

We hope to recover edges with high commonalities and those which consistently rank highly across the remaining cell types where they didn’t make to the top 5% correlations to derive a core network. Similar to the analysis of the single-cell brain dataset in the main text, we used a rank aggregation analysis to inform this edge commonality cutoff. Using various edge commonality groups covering its whole spectrum, per edge commonality group, we randomly sampled 100 edges if this group contained at least 100 edges, otherwise use all available edges. For each edge, we first collected the cell types that this edge was absent (didn’t make to the top 5% correlations), and then calculated the adjusted P value on these cell types using a rank aggregation method. The result showed that the negative log-transformed adjusted P value distributions increase together with edge commonalities, similar to the pattern observed in the single-cell fly brain atlas data, but much less pronounced (**Fig. S9D**). In particular, all edges but one edge from edge commonality group 37 sampled from the edge commonality groups ≥ 35 were highly ranked in the remaining cell types (P < 0.05), whereas only ≤ 1% of the tested edges shared by up to 11 cell types (edge commonality ≤ 11) were significant (P > 0.05). Based on this rank aggregation analysis, we chose edge commonality 35 as the cutoff to select core network edges, which resulted in a core network in the *Drosophila* head with 20 genes connected by 21 edges (**Fig S9E)**. We collected all 21 edges and showed their ranks across cell types. Similar to the brain-cell core network in the main text, the head-cell network lacked edges found in all cells, but its edges were highly ranked in the remaining cell types (**Fig S9F**). Compared with the core network which composes 205 genes in the single-cell fly brain data, 7 genes overlapped. Simulations showed significant overlap between core networks inferred from the two datasets (empirical P value = 0.00099, calculated as (n+1)/(N+1), where n indicates the number of permutations that are larger than the observed number and N the total number of permutations, **Fig S9G)**, validating our discovery in the main text.

**References**

Baker BM, Mokashi SS, Shankar V, Hatfield JS, Hannah RC, Mackay TFC, Anholt RRH. 2021. The Drosophila brain on cocaine at single-cell resolution. *Genome research* **31**: 1927–1937.

Bozhilova L v, Pardo-Diaz J, Reinert G, Deane CM. 2021. COGENT: evaluating the consistency of gene co-expression networks. *Bioinformatics* **37**: 1928–1929.

Iacono G, Massoni-Badosa R, Heyn H. 2019. Single-cell transcriptomics unveils gene regulatory network plasticity. *Genome biology* **20**: 1–20.

Li H, Janssens J, de Waegeneer M, Kolluru SS, Davie K, Gardeux V, Saelens W, David F, Brbić M, Leskovec J. 2021. Fly Cell Atlas: a single-cell transcriptomic atlas of the adult fruit fly. *bioRxiv*.
